# Spatiotemporal lineage tracing reveals the dynamic spatial architecture of tumour growth and metastasis

**DOI:** 10.1101/2024.10.21.619529

**Authors:** Matthew G. Jones, Dawei Sun, Kyung Hoi (Joseph) Min, William N. Colgan, Haoyu Wang, Tivadar Török, Erik C. Cardoso, Luyi Tian, Jackson A. Weir, Victor Z. Chen, Luke W. Koblan, Kathryn E. Yost, Nicolas Mathey-Andrews, Edridge D’Souza, Andrew J.C. Russell, Robert R. Stickels, Karol S. Balderrama, William M. Rideout, Min Dai, Giovanni Marrero, Vipin Kumar, Anjali Saqi, Benjamin Herzberg, Benjamin Izar, Howard Y. Chang, Joo-Hyeon Lee, Tyler Jacks, Fei Chen, Jonathan S. Weissman, Nir Yosef, Dian Yang

## Abstract

Tumour progression is driven by dynamic interactions between cancer cells and their surrounding microenvironment. Investigating the spatiotemporal evolution of tumours can provide crucial insights into how intrinsic changes within cancer cells and extrinsic alterations in the microenvironment cooperate to drive different stages of tumour progression. Here, we integrate high-resolution spatial transcriptomics and evolving lineage tracing technologies to elucidate how tumour expansion, plasticity, and metastasis co-evolve with microenvironmental remodelling in a *Kras;p53*-driven mouse model of lung adenocarcinoma. We find that rapid subclonal expansion contributes to a hypoxic, immunosuppressive, and fibrotic microenvironment that is associated with the emergence of pro-metastatic cancer cell states. Furthermore, metastases arise from spatially-confined subclones of primary tumours and remodel the distant metastatic niche into a fibrotic, collagen-rich microenvironment. Together, we present a comprehensive dataset integrating spatial assays and lineage tracing to elucidate how sequential changes in cancer cell state and microenvironmental structures cooperate to promote tumour progression.

## MAIN

Tumour progression is driven by the dynamic interactions between cancer cells^1,2^ and the their surrounding microenvironment^3,4^. In this process, as cancer cells accumulate genetic and epigenetic alterations, the microenvironment exerts selective pressures through factors such as spatial constraints^5,6^, signalling molecules^7^, nutrient and oxygen availability^8,9^, and immune infiltration^3,10^ among other phenomena. In turn, tumour growth remodels the surrounding microenvironment, for example, by restructuring the extracellular matrix and altering the composition and state of infiltrating stromal cells^11^. Systematically characterizing the cell intrinsic and extrinsic effects that drive tumour subclonal selection, cellular plasticity, and metastasis will not only provide insights into the principles of tumour evolution but also carry clinical implications. To accomplish this, one must study a tumour’s evolutionary dynamics alongside its microenvironmental composition in the native spatial context.

Integrating tumour phylogenetic analysis, the study of lineage relationships of cancer cells within a tumour^12–17^, with spatial information provides a comprehensive framework for understanding the interplay between tumour microenvironment and progression. Specifically, spatially resolved phylogenetic studies enable one to approach key questions in cancer evolution such as, what are the major spatial communities that exist in tumours, and how do these relate to tumour stage? From which spatial niches do subclonal expansions arise during tumour progression, and how does this relate to tumour plasticity and the capacity to seed metastases? And, how does the spatial growth pattern of tumour progression shape the surrounding microenvironment? Early studies reconstructing tumour phylogenies from multi-region sampling of patient tumours uncovered the spatial heterogeneity of genetic changes within tumours and have demonstrated the dynamics of tumour growth and spatially-constrained origins of metastatic dissemination^18–24^. More recently, spatial genomics approaches have further elucidated how the spatial distribution of genome alterations leads to clonal outgrowth, dispersion of subclones with distinct driver mutations, interactions with the immune system, and metastasis^25–29^. While these studies have greatly enhanced our understanding of how tumours grow in space and time, they can be limited in their ability to either resolve high-resolution spatial organisation, infer deeper phylogenetic relationships of cancer cells, or simultaneously measure the microenvironmental composition and gene expression.

The development of molecular recording technologies that install evolving lineage-tracing barcodes^30–40^ and associated computational tools^41–46^ enable the reconstruction of high-resolution phylogenies for studying tumour evolution^13^. Typically, these lineage-tracing technologies employ genome-editing tools, such as CRISPR/Cas9, to introduce heritable and irreversible mutations progressively at defined genomic loci, which can be transcribed and thus profiled with single-cell RNA-seq. In cancer, initial studies applied this technology to track the metastatic dynamics of cancer cell lines transplanted into mice^47–49^. Previously, we described a lineage-tracing enabled genetically-engineered mouse model of *Kras^LSL-G12D/+^;Trp53^fl/fl^-*driven lung adenocarcinoma (KP-Tracer) to continuously track tumour evolution from nascent transformation of single cells to aggressive metastasis^50^. In this system, intratracheal delivery of *Cre* recombinase using viral vectors simultaneously induces Cas9-based lineage tracing and tumour initiation. This model recapitulates the major steps of the evolution of human lung adenocarcinoma, both molecularly and histopathologically^51–55^. Using this system, we recently identified subclonal expansions, quantified tumour plasticity, traced metastatic origins and routes, and disentangled the effect of genetic drivers on tumour evolution. However, as our previous applications have relied on studying dissociated single cells, it has remained unclear how key tumour evolutionary properties are associated with microenvironmental changes.

Here, we present an integrated lineage and spatial platform for tracking tumour evolution *in situ* by applying high-resolution spatial transcriptomics to our lineage tracing-enabled KP-Tracer model. Using two complementary spatial transcriptomics assays – Slide-seq^56,57^ with spot-based coverage at 10𝜇m near-cell resolution of large tissue fields-of-view, and Slide-tags^58^ with higher molecular sensitivity and spatial profiling of individual nuclei – we produce a comprehensive spatial transcriptomics dataset of *Kras;p53-*driven lung adenocarcinoma evolution. Integrating these spatial transcriptomics data with inferred cancer cell lineages uncovered robust spatial communities associated with tumour progression, including the formation of a hypoxic tumour interior during rapid tumour subclonal expansion. Our analysis additionally reveals that this hypoxic environment is associated with pervasive tissue remodelling characterised by fibrosis, priming of immunosuppressive immune cells and rewiring of cellular interactions, and the emergence of a pro-metastatic epithelial-to-mesenchymal transition (EMT). Together, this study provides a scalable platform for studying the relationship between tissue architecture and tumour progression, revealing key insights into the ecological and evolutionary dynamics underpinning tumour evolution at unprecedented resolution.

## RESULTS

### An integrated lineage and spatial platform for studying tumour evolution

To study tumour evolution while preserving the native spatial context of cancerous and stromal tissue, we integrated spatial transcriptomics methods with Cas9-based lineage-tracing technology in our previously described KP-Tracer model of lung adenocarcinoma^50^. This model is built upon the well-characterised model of *Kras;Trp53*-driven lung adenocarcinoma^51,52,54,55^ and is equipped with a *Cre-*inducible Cas9-based evolving lineage tracer that is able to continuously record high-resolution cell lineages over months-long timescales^32,41^. Introduction of *Cre* into individual lung cells in the adult animal both induces the oncogene mutations (i.e., expression of *Kras^G12D^* and homozygous loss of *p53*) and initiates Cas9 expression. Cas9 then introduces irreversible and heritable insertions and deletions (“indels”) at defined genomic “target sites”, each discernable by a random 14bp integration barcode (“intBC”) and expressed as a polyadenylated transcript. As most sequencing-based spatial transcriptomics assays capture polyadenylated transcripts from tissue sections^56–60^, applying these assays to the KP-Tracer model yields simultaneous measurement of spatially-resolved cell transcriptional states and lineage relationships.

We initiated lung tumours and lineage-tracing in alveolar type II (AT2) cells (a major cell of origin for lung adenocarcinoma) by intratracheally delivering adenovirus expressing *Cre* recombinase under the control of an AT2 cell-specific, surfactant Protein C (SPC) gene promoter^61^. Twelve to sixteen weeks post tumour initiation, tumour bearing lungs were harvested for cryopreservation, and then sectioned and applied to spatial transcriptomics arrays (**Fig. 1a; Methods**). To comprehensively profile the spatiotemporal evolution of tumour progression, we utilised two complementary spatial transcriptomics technologies: Slide-seq^56,57^ that captures transcriptomic states of “spots” at near-cellular 10𝜇m resolution in continuous, large fields-of-view (up to 1cm x 1cm); and Slide-tags^58^ that sparsely samples individual nuclei for transcriptomic profiling and provides accurate spatial localisation for a subset of these nuclei (typically ∼50-70%). Together, this combination marries the scale of Slide-seq and true single-nucleus resolution of Slide-tags to jointly measure spatially resolved cell lineage and unbiased transcriptomic states in the native tumour microenvironment.

**Figure 1.**
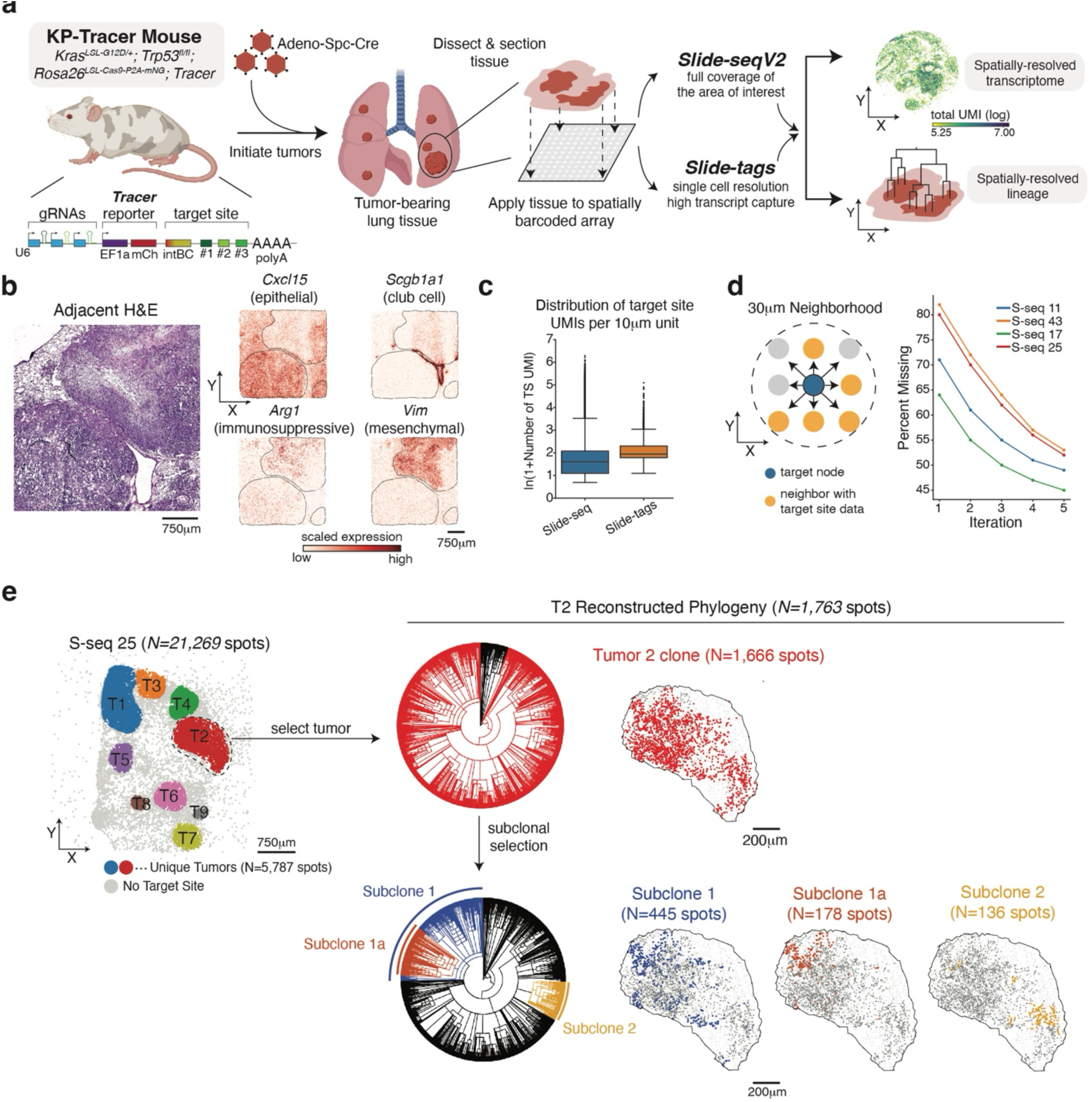
An integrated lineage and spatial platform enables high-resolution analysis of tumour evolution *in vivo*. **(a)** Schematic of experimental workflow for integrated, spatially resolved lineage and cell state analysis. In KP-tracer mice, oncogenic *Kras^G12D/+^*;*Trp53^-/-^* mutations and Cas9-based lineage tracing were simultaneously activated upon administration of adenovirus carrying SPC promoter-driven Cre recombinase. After 12-16 weeks, mice were sacrificed, and cryopreserved tumour-bearing lungs were sectioned for spatial profiling with Slide-seq and Slide-tags technologies. Libraries were prepared and sequenced to study spatially resolved lineages and transcriptional patterns. S-seq 30 is used as a representative example for total UMI capture in a spatial array. Biorender was used to create parts of this schematic. **(b)** Representative H&E staining and spatially resolved gene expression data for a lung section carrying three tumours (black line). Log-normalised, scaled counts for epithelial-like (*Cxcl15* and *Scgb1a1*), immunosuppressive myeloid (*Arg1*), and mesenchymal cells (*Vim*) are shown. **(c)** Distribution of the number of target-site UMIs for Slide-seq and Slide-tags data. Ln(1+x) counts are shown. Boxplots show the quartiles of the distribution, and whiskers extend to 1.5x the interquartile range. **(d)** Schematic of spatial imputation of lineage-tracing data in 30𝜇𝑚 neighbourhoods (left) and representative examples of missingness left after each of 5 iterations of spatial imputation. **(e)** Representative spatially resolved lineages in spatial array S-seq 25 profiling a lung section carrying 9 distinct tumours. Reconstructed lineages are displayed for a representative tumour, T2. Successive nested subclones displaying both shared and distinct lineage states in unique colors are indicated on the phylogenetic tree and mapped spatially. Lineages marked in black spots not included in the designated subclone. Overall, spots that are more related in lineage tend to be spatially coherent.

With these two technologies, we comprehensively profiled tumour-bearing lungs across various stages of progression with 44 Slide-seq arrays and 5 Slide-tags arrays (**Extended Data Fig. 1a-c; Methods**; **Supplementary Table 1**). The resulting datasets provided spatial profiling of distinct domains in tumour-bearing tissues characterised by the expression of canonical marker genes and corroborated by paired H&E: for example, in the tumour-bearing lung we found that *Cxcl15* and *Scgb1a1* marked epithelial-like domains, representing alveolar and club cells, respectively. Moreover, histologically aggressive regions were marked by *Vim* (characteristic of mesenchymal-like cancer cells) and *Arg1* (characteristic of immunosuppressive myeloid cells^62^) **(Fig. 1b**). Altogether, these datasets provide high-resolution views into the microenvironmental context and organisation of tumours.

### Computational tools enable the inference of spatially resolved cancer cell phylogenies

As the KP-Tracer system expresses lineage tracing target-sites as poly-adenylated transcripts, we next turned to evaluating the recovery of these target sites from the complementary spatial transcriptomics platforms. Reassuringly, we detected target-site transcripts robustly across tens-of-thousands of spots or nuclei in these spatial datasets, with Slide-tags data having more consistent detection of target-sites as expected (**Fig. 1c; Extended Data Fig. 1d-e**).

While Slide-tags provided true single-cell measurements and thus were amenable to previously-described lineage reconstruction approaches^41,44^, there were two predominant analytical challenges in reconstructing tumour phylogenies of tens-of-thousands of spots observed in Slide-seq data. First, Slide-seq captures RNA molecules with near-cellular resolution, meaning that each spot may contain RNAs originating from multiple cells^57^; similarly, cells with distinct lineage states can be captured in a single spot, which we term “conflicting states”. As prior phylogenetic reconstruction algorithms for Cas9-lineage tracing data presume mapping of cells to single states, we first implemented new Cassiopeia-Greedy^41^ and Neighbour-Joining^63^ variants that could use many conflicting states during reconstruction (**Methods**). We also tested the effects of three strategies for preprocessing conflicting states via simulation: (1) a strategy that used all conflicting states observed in a spot along with the abundance of each state in that spot (“all states”); (2) all conflicting states observed in a spot, but without considering their abundance (“collapse duplicates”); or (3) a strategy that used only the most abundant state (“most abundant”). We found that the second strategy (“collapse duplicates”) performed most robustly (**Extended Data Fig. 2a**; **Methods**).

A second challenge is that Slide-seq assays (and to a lesser extent Slide-tags) have an increased missing data rate relative to droplet-based single-cell assays^64^. As expected, we observed overall lower target-site transcript capture (and thus higher missing data) in Slide-seq datasets (**Extended Data Fig. 1d and Extended Data Fig. 2b**). We hypothesised that spatial relationships could be used to overcome this sparsity, which was supported by our observations that indel states were coherent within small spatial neighbourhoods (**Extended Data Fig. 2c-d**). We therefore developed an inferential approach that predicted missing lineage-tracing states from spatial neighbours (within 30 𝜇 m of a target node) with sufficient recovery (at least 3 UMI supporting a target site intBC-indel combination; **Fig. 1d**). We first tested the feasibility of this approach using simulations of lineage tracing data on spatial arrays using Cassiopeia (**Methods)**. We found that missing lineage-tracing barcodes could consistently be recovered at high accuracy (**Extended Data Fig. 2e**), and that spatial imputation followed by tree inference by a hybrid algorithm consisting of the Cassiopeia-Greedy and Neighbour-Joining algorithms resulted in the best reconstructions, especially in high-dropout regimes (**Extended Data Fig. 2f-g**; **Methods**). Next, we tested our ability to recover held-out target site data from real Slide-seq data and similarly found that missing data could be robustly recovered by spatial predictions, resulting in a median accuracy of 90% on imputing held-out data across all experiments, matching our simulation results (random predictions had a median accuracy of 67% and yielded 29% fewer imputations; **Extended Data Fig. 2h**). As expected, more frequent alleles had higher imputation accuracy (**Extended Data Fig. 2i**; **Methods**). Over multiple iterations of this imputation algorithm, we found that we could recover up to 58% of missing data (4-58%, on average 31% across datasets), resulting in comparable missing data rates to previous reports using single-cell approaches that have enabled robust tree reconstruction and biological insights (**Fig. 1d, Extended Data Fig. 2j**). Though we only retain high-confidence imputations, and our benchmarks point to the promise of this spatial imputation in this context, there are notable caveats especially in the case of cell migration (see **Discussion**). Combining Slide-seq data and validation from orthogonal trees provided by Slide-tags establish a foundation for studying the spatial lineages of cancer cells.

Together, these computational improvements enabled us to build lineages of cancer cells in the native context of a tumour’s microenvironment at unprecedented resolution (**Fig. 1e**). Our lineages revealed phylogenetic relationships in structured spatial environments and enabled us to explore the spatial localisation of increasingly related subclones within the same tumour (**Fig. 1e**, **Extended Data Fig. 2k)**. Furthermore, in comparing trees built from Slide-tags data, we observed copy-number variation (CNVs) largely corroborated lineages (as previously shown^50^), though there were far fewer CNVs per cell than lineage states generated by the lineage tracer (**Extended Data Fig. 3**). With these data and approaches, we turned to investigating the relationship between changes to the microenvironmental architecture and tumour progression.

### Spatial transcriptomics reveal the ecosystems of lung adenocarcinoma

While recent efforts have studied the composition of tumours in this model using single-cell approaches^50,54,55^, it has remained challenging to profile the spatial organisation of these cell types. To address this, we leveraged the complementary insights gained from the high sensitivity, true single-nucleus measurements of Slide-tags and the broad field-of-view of Slide-seq to perform a systematic analysis of tumour spatial organisation across stages of progression observed in our 49 spatial transcriptomics arrays representing more than 100 tumours.

Focusing first on the true single nuclei profiled with Slide-tags, we performed fine-grained annotation of clusters consisting of normal epithelial, stromal, immune, and tumour cells (determined by canonical marker genes and the presence of active lineage-tracing edits) (**Fig. 2a-b**; **Extended Data Fig. 4a**; **Methods**). In addition to annotating previously described tumour and normal epithelial cells in this model^50,55^, we identified a previously undescribed tumour cell state characterised by the expression of neuronal genes such *Piezo2* and *Robo1,* the endothelial marker *Pecam1,* maintenance of the lung-lineage transcription factor *Nkx2-1*, and absence of *Vim* (**Extended Data Fig. 4b-c**). Although this cell type expressed active lineage tracing marks in our system, it is likely that this cell type was excluded in previous studies^50,55,65^ by purifying cancer cells against CD31 expression (also known as *Pecam1*, expressed in this population) prior to transcriptomic profiling; this highlights the advantage of spatial transcriptomics in profiling all cells and communities, eliminating potential biases arising from tissue dissociation and preparation. In the immune and stromal compartment, we observed large macrophage, fibroblast, and endothelial populations with lower representation of B cells and dendritic cells (**Fig. 2a**; **Extended Data Fig. 4a**). Among macrophages, we detected *SiglecF+* tissue-resident alveolar macrophages and three distinct tumour-associated macrophage (TAM) populations: *Vegfa+* TAMs, immunosuppressive *Arg1+* TAMs, and proangiogenic *Pecam1+* TAMs (**Fig. 2a)**. We additionally detected a diverse set of cancer-associated fibroblasts (CAFs): a mesothelial-like *Wt1+* population, an inflammatory-like CAF (“iCAF”) population expressing the complement gene *C7* and *Abca8a*, and a myofibroblast-like CAF (“myCAF”) population expressing *Postn* (**Fig. 2a, Extended Data Fig. 4a**).

**Figure 2.**
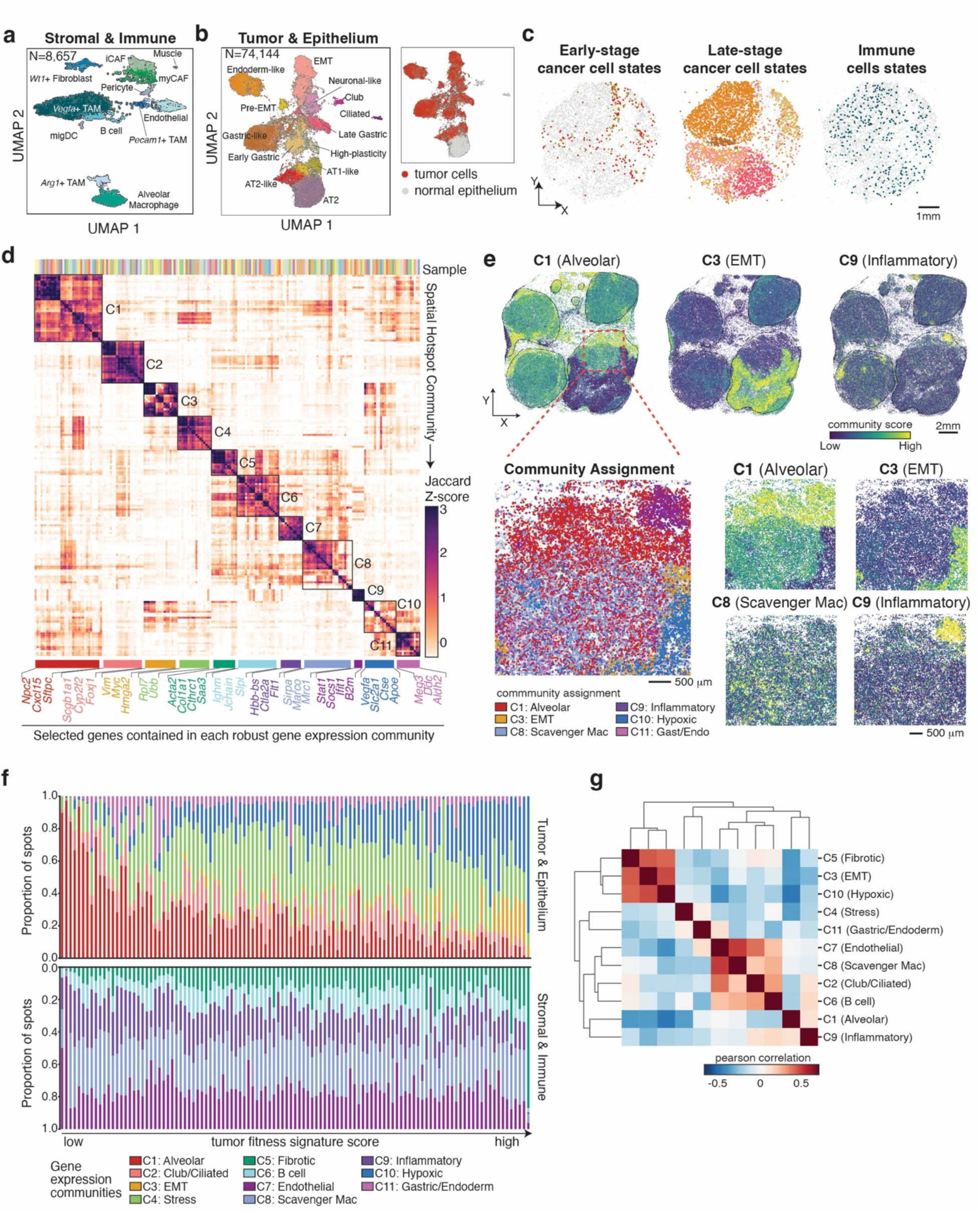
Diverse spatial gene expression communities emerge during KP-tracer tumour progression. **(a-b)** UMAP projections of Slide-tags data on tumour bearing lungs from KP-Tracer mice, annotated by cell type. **(a)** Slide-tags data corresponding to all stromal and immune cell types: *Cd45+* immune cells and other non-epithelial stromal cells. **(b)** Slide-tags data corresponding to all cancer and normal epithelial cells. Inset indicates where cancer cells are found in this projection. **(c)** Representative spatial projections of early-stage and late-stage cancer cell states, and immune cell types from Slide-tags analysis of KP-Tracer tumour bearing lung (shown on S-tags 3). Colors correspond to those in UMAP projections in **(a-b)**. **(d)** Heatmap of Z-scored Jaccard overlap between genes contained in spatial gene expression communities. Each row or column is a community, defined as a set of spatially autocorrelated genes identified with Hotspot, and robust spatial gene expression communities are determined by hierarchical clustering and indicated by annotated blocks. The Slide-seq sample from which a community is identified is indicated by unique colors on the top of the heatmap. Representative genes specific to each spatial community are highlighted at the bottom of the heatmap. **(e)** Community scores of selected spatial communities projected onto a representative Slide-seq dataset of a tumour bearing lung with 4 major tumours (S-seq 43). Tumour boundaries are indicated with black lines (top). Zoom in of region showing community assignments and scores for a selection of communities (bottom). **(f)** Proportion of gene expression community assignments across all KP lung tumours in the Slide-seq dataset, ordered by increasing fitness signature scores. Each bar indicates a single segmented tumour in the Slide-seq dataset. Top: communities that are more related to tumour or epithelial programs. Bottom: communities that are related to stromal and immune programs. **(g)** Heatmap reporting Pearson correlation of community abundances across all tumours in the Slide-seq data.

To explore the spatial localisation of these diverse cell states, we assigned spatial locations to Slide-tags nuclei and spatially projected cell identities (**Methods**). Consistent with previous characterisations of Slide-tags spatial mapping rates^14^, we found that approximately 50% of nuclei could be confidently assigned to a spatial location (**Extended Data Fig. 4d**). Across the five Slide-tags arrays, we observed a distinct pattern where less aggressive, “early-stage” tumour cell states (i.e., AT2- and AT1-like cancer cells, indicated by expression of active lineage marks and distinct gene expression from normal AT2 and AT1 cells) co-localized on the periphery of tumour sections consisting of more aggressive “late stage” tumour cells (**Fig. 2c, Extended Data Fig. 4e**). Similar to previous work in this model^66^, we also found that distinct immune and stromal cell types exhibited differential infiltration – for example, Alveolar Macrophages and iCAFs were typically found outside tumours, whereas *Arg1+* TAMs and myCAFs were more likely to be found within tumours (**Fig. 2c, Extended Data Fig. 4e)**.

The spatially-localised transcriptional signatures observed with Slide-tags motivated us to pair this approach with Slide-seq assays to survey the spatial gene expression communities across large tissue areas in tumours. We thus turned to the 44 tissue sections assayed with Slide-seq that collectively represent more than 100 tumours at various tumour stages. To identify modules of genes that were recurrently spatially co-expressed across multiple samples, we employed the Hotspot^33^ algorithm (**Methods**). Our analysis revealed 11 recurrent spatial gene modules, hereafter referred to as “communities” (**Fig. 2d-e; Supplementary Table 2**), that we annotated by inspecting the genes contained within communities and evaluating the expression level of community genes (captured in a “community score”) in cell types identified by Slide-tags data (**Extended Data Fig. 4f; Extended Data Fig. 5a)**.

The genes contained within these transcriptional communities represent a variety of co-localised gene expression states: for example, an early-stage alveolar-like community contained genes marking epithelial cells such as *Sftpc* and *Cxcl15* (“C1: Alveolar”), a hypoxic community contained canonical marker genes of hypoxia such as *Slc2a1* (also known as *Glut1*) (“C10: Hypoxia”), and an epithelial-to-mesenchymal (EMT) community contained genes such as *Vim*, up-regulation of *Myc* signalling, and metastasis-related genes such as *Hmga2* (“C3: EMT”; **Fig. 2d-e, Extended Data Fig. 5a**). In addition to fibroblast (C5), B cell (C6), and endothelial (C7) communities, we identified two distinct immunoregulatory-related communities. The first community contained genes associated with scavenger-like macrophages like *Marco* and *Mrc1* (“C8: Scavenger Mac”); a second community contained genes characteristic of inflammation such as *B2m*, *Stat1*, and *Ifit1* (“C9: Inflammatory”). As these communities describe genes co-expressed in spatial proximity, they provide insights into possible intercellular interactions. For example, the EMT and hypoxic communities (C3 and C10) contained genes associated with macrophage recruitment (e.g. *Csf1)* and polarisation to immunosuppressive states that have been previously reported to promote aggressive cancer phenotypes (e.g., *Arg1*^62^ and *Spp1*^67^), while the Inflammatory community (C9) contained *Cxcl9* that has been previously reported in anti-tumour macrophage polarisation^67^ (**Extended Data Fig. 5a**).

To inspect the distribution of these communities across large tissue sections profiled with Slide-seq, we quantified community scores for each spot and assigned spots to the community with the highest score (**Fig. 2e-f, Extended Data Fig. 5b-c**). In comparing histology from an adjacent layer to the community scores, we found co-localisation between areas indicating high tumour grade (as indicated by histology) and high scores for EMT, hypoxic, and fibrotic communities (C3, C10, C5; **Extended Data Fig. 5b**). We next asked how the distribution of community assignments varied over tumour stages using a gene set signature we previously identified to robustly associate with tumour progression (termed a “fitness signature”)^50^ (**Fig. 2f; Extended Data Fig. 5c**; **Methods**). Specifically, this fitness signature contains genes that are associated with subclonal expansions in this model, and their collective activity (i.e., “score”) reflects tumour progression towards an aggressive, pro-metastatic state. Consistent with the definition of this signature, after ranking tumours by their fitness signature score and inspecting the proportion of community assignments, we observed that early-stage tumours were dominated by epithelial, endothelial, and inflammatory communities (C1, C7, and C8, respectively) but that late-stage tumours had larger fractions of EMT, hypoxic, and fibroblast communities (C3, C10, and C5, respectively; **Fig. 2f, Extended Data Fig. 5c**). Moreover, we found that overall abundances of EMT, hypoxic, and fibroblast community assignments (C3, C10, and C5, respectively) were correlated across all tumours; conversely, they were anticorrelated with the abundances of alveolar and inflammatory communities (C1 and C8, respectively) (**Fig. 2g**).

Together, these analyses unite the unique advantages of Slide-tags and Slide-seq assays to provide a consensus set of spatial communities that highlight differential immune and stromal activation and localization patterns across tumour progression in KP tumours. These observations motivated us to next integrate our phylogenies to understand how the spatiotemporal dynamics of these communities are associated with tumour plasticity and subclonal expansion.

### Rapid tumour subclonal expansion contributes to a hypoxic niche with decreased cancer cell plasticity

Integrating cell state information with high-resolution phylogenies can offer new insights into various aspects of tumour evolution. In our previous work, we introduced frameworks to quantify the historical record of subclonal growth rates (“phylogenetic fitness”) and the number of cell-state transitions in each cell’s lineage (“single-cell clonal plasticity”) directly from phylogenies^50,68^. We described a model whereby KP-Tracer tumour progression is driven by the loss of an initial AT2-like cell state and accompanying increases in single-cell clonal plasticity and transcriptional heterogeneity; in turn, these high-plasticity cells provide a diverse pool of transcriptional states from which high-fitness, low-plasticity subclones with increased metastatic ability and expression for EMT markers like *Vim* and *Hmga2* are selected^50^. Consistent with this previous work, the tumours studied with this spatial-lineage platform showed an overall distribution where transient increases in plasticity are followed by the selection of low-plasticity, high fitness subclones (**Extended Data Fig. 6a**). Since these measures of fitness and clonal plasticity are computed from phylogenies (and not cell-intrinsic features), we reasoned that we could extend our previously reported model by relating these phylodynamic changes to the surrounding microenvironment, thereby identifying cancer cell extrinsic factors associated with tumour fitness and plasticity.

As the measurement of phylogenetic fitness reports on the history of subclonal growth, spatially-resolved phylogenies are well suited to understanding the relationship between tumour growth patterns and microenvironmental properties^22,69^. In one representative Slide-seq example (S-seq 40), we found an expanding subclone with high phylogenetic fitness localised to a tumour interior characterised by late-stage Hypoxic and EMT communities (C10 & C3) while the tumour periphery had lower phylogenetic fitness and was marked by the Alveolar community (C1) (**Fig. 3a**). This co-localisation of high phylogenetic fitness with hypoxic regions was supported by four lines of evidence: first, we found that phylogenetic fitness was correlated with the orthogonal, previously-described fitness signature^50^ (Pearson’s *r* = 0.4; **Extended Data Fig. 6b**). Second, the emergence of a large CNV-defined subclone (containing an amplification of the *Kras* locus on chromosome 6) co-localized with the hypoxic regions (**Extended Data Fig. 6c**). Third, in a systematic analysis of all Slide-seq tumours, we found that the EMT and Hypoxic communities were most strongly correlated with phylogenetic fitness (**Extended Data Fig. 6d**). Finally, across all high-resolution Slide-tags arrays, we similarly found that the late-stage states (e.g., EMT) were most likely to be found in regions that had previously undergone subclonal expansion (**Extended Data Fig. 6e**). These orthogonal data collectively support the observation that the co-localisation of expansion and hypoxia is consistent across tumours and is not an artifact of tree reconstruction or the near-cell resolution of Slide-seq.

**Figure 3.**
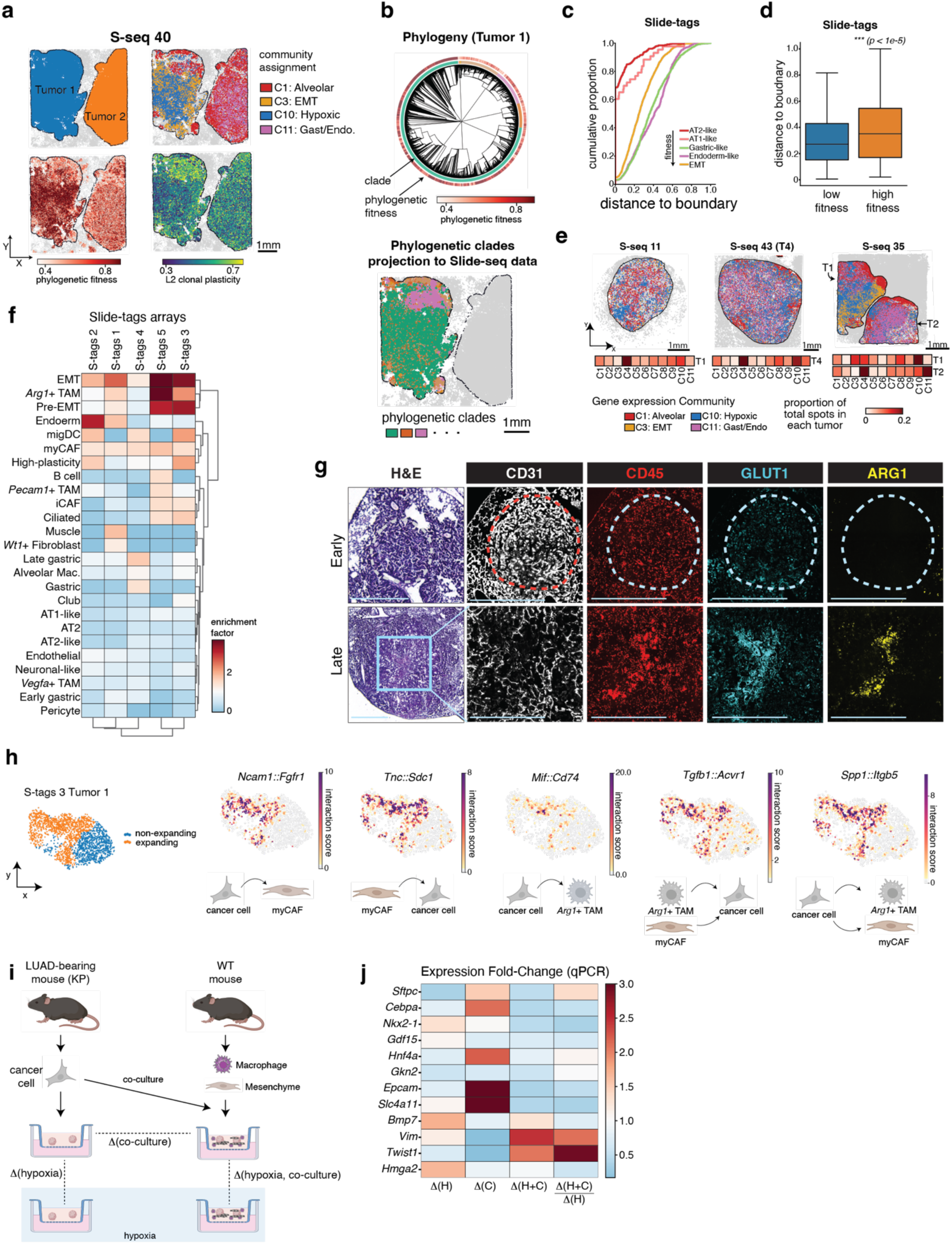
Subclonal expansions associate with microenvironmental remodelling towards a hypoxic, fibrotic, and immunosuppressive state. **(a)** A representative Slide-seq array containing two tumours (S-seq 40) is shown with spatial projections of tumour annotations, selected gene expression community assignments, phylogenetic fitness, and L2 clonal plasticity. **(b)** Reconstructed phylogeny and spatial localization of phylogenetic subclades for Tumour 1 from the representative Slide-seq dataset (S-seq 40) example shown in **(a)**. The phylogeny is annotated by subclonal clade assignment (inner colour track) and phylogenetic fitness (outer colour track). **(c)** Cumulative density distributions for normalised Euclidean distance to nearest non-tumour cell (i.e., tumour boundary) for five selected major cancer cell states across all Slide-tags arrays. Cancer cells in high-fitness-associated cell states (e.g. EMT, Endoderm-like, Gastric-like) locate further away from the tumour boundary than those in low-fitness-associated states (AT2-like, AT1-like). Distance is normalised to unit scale (0-1). **(d)** Distribution of normalised Euclidean distances to nearest non-tumour cell (i.e., tumour boundary) for high-fitness and low-fitness cells (defined here as having phylogenetic fitness greater than the 90^th^ or less than the 10^th^ percentiles, respectively). High-fitness cells are significantly further away from the tumour boundary (*p<1e-5*, wilcoxon rank-sums test). Boxplots show the quartiles of the distribution, and whiskers extend to 1.5x the interquartile range. **(e)** Representative Slide-seq examples showing the evolution of the spatial gene expression communities following tumour progression (left to right). Selected community assignments are displayed, and full proportion of assignments are reported in 1D heatmaps under each spatial dataset. **(f)** Clustered heatmap of enrichments of cell type abundances in spatial neighbourhoods of high- and low-fitness cells in 5 Slide-tags arrays. Values > 1 indicate that a cell type is more abundant (i.e., enriched) in neighbourhoods of cells with high fitness. Cell type names are identical to those reported in Fig. 2a**-b**. **(g)** H&E and paired immunofluorescence staining of endothelial-cell marker CD31, immune cell marker CD45, hypoxia-reporter GLUT1, and immunosuppressive myeloid marker ARG1 in representative KP tumours. The interior of large, late-stage tumours is marked with a decrease of endothelial cells (CD31) and increases of hypoxia (GLUT1) and immunosuppressive myeloid cells (ARG1, CD45). Scale bars = 1mm. **(h)** Representative examples of expansion-specific ligand-receptor interaction pairs. Interaction pair is indicated by (ligand::receptor) and cell types and directionality of interaction are indicated below by arrows and cell types. Values are interaction scores based on the expression of the ligand-receptor pair in the indicated spot. **(i-j**) Co-culture of KP tumour organoids, alveolar macrophages, and lung mesenchyme in hypoxic conditions induce a stronger EMT response in cancer cells than hypoxia alone. (i) Experimental setup: cancer cells were sorted from KP tumours and alveolar macrophages and lung mesenchymal cells were sorted from normal mice. The 3D primary organoids were derived from cancer cells and were cultured individually or in co-culture in normal and hypoxic conditions. (j) Gene expression fold-changes across conditions, measured by qPCR. (H: Hypoxia; C: Co-culture)

The localisation of expanding subclones characterised by aggressive gene expression states in a representative Slide-seq example (S-seq 40) prompted us to hypothesise that rapid subclonal expansions may create a layered architecture whereby expanding subclones dominate a central region surrounded by non-expanding cells (**Fig. 3a-b**). Focusing first on this representative Slide-seq example, we observed that multiple low-fitness areas of Tumour 1 could be grouped together in a phylogenetic subclade despite being geographically distant (though many indels were shared across the tree, these low-fitness, distant cells were marked by the shared absence of indels marking the expanding region) (**Fig. 3a-b**; **Extended Data Fig. 6f**). Though this pattern could be generated many ways (e.g., independent migration of several subclones), the most parsimonious interpretation suggests that these scattered low-fitness cells were in close spatial proximity during the early stage of tumour growth but were later pushed to the tumour periphery because of a subclonal expansion event. To investigate the consistency of this phenomenon, we next quantified the phylogenetic fitness of individual cancer cells derived from high-resolution Slide-tags arrays on multiple tumours and inspected the spatial distribution of subclonal expansion. In this analysis, we also found that the tumour interior in Slide-tags data was more likely to contain cells with more aggressive gene expression states (e.g., Endoderm-like and EMT states) and higher phylogenetic fitness as inferred from reconstructed trees (**Fig. 3c-d** *p < 1e-5*, wilcoxon rank-sums test; **Fig. 2c**; **Extended Data Fig. 4e**).

Recent work in this model has underscored the importance of epigenetic remodelling during tumour progression, often manifesting as rapid cell-type transitions that can be quantified using a phylogeny-based single-cell clonal plasticity measure^50,54^. Motivated by the observed interplay between expansion and microenvironmental remodelling, we next turned to investigating the association between spatial features and changes in single-cell clonal plasticity. Starting in the representative Slide-seq example (S-seq 40), we observed that low-plasticity clones in Tumour 1 co-localised with high-fitness regions in the tumour interior whereas the high-plasticity regions of Tumour 2 (which lacked a subclonal expansion) appeared to lack spatial organisation (**Fig. 3a**). Consistent with this, we found that the high-fitness Hypoxic and EMT communities, and related states, were associated with lower plasticity across all Slide-seq and Slide-tags datasets (**Extended Data Fig. 6g-h**). To better understand how transient increases in plasticity associated with the subclonal expansions observed across Slide-seq datasets (**Extended Data Fig. 6a**), we further examined the transition to subclonal expansion in arrays profiled with Slide-tags (**Extended Data Fig. 6i-k**). Across our Slide-tags data, we found there was little spatial organisation of high-plasticity cells in tumours without detectable subclonal expansion (as measured by Moran’s *I* autocorrelation statistic^70^), whereas low-plasticity cells were spatially localized to the tumour centre in tumours after expansion (**Extended Data Fig. 6j-k**; **Methods**).

Collectively, these data support a model whereby the tumour microenvironment is sequentially remodelled by subclonal expansion, culminating in an interior hypoxic region and eventually the emergence of a late-stage, pro-metastatic EMT state. Moreover, our analysis of clonal transcriptional plasticity refines our previous model of tumour progression: after loss of the initial AT2-like identity and an increase in clonal plasticity, rapid subclonal expansion drives spatial reorganization of the tumour and leads to the formation of a hypoxic region, which may in turn induce and stabilize the EMT of cancer cells. As evidenced by examples of tumours across various stages, this model is characterised by the exclusion of early-stage communities (e.g., C1: Alveolar) to the tumour periphery while subclonal expansions contribute to the acquisition of a low-plasticity, high-fitness Hypoxic community (C10) and eventual transition to an EMT community (C3) (**Fig. 3e**; **Fig. 2f**; **Extended Data Fig. 6i**).

### Subclonal expansion is accompanied by immunosuppressive and fibrotic microenvironmental remodelling

As our Slide-seq data suggest that the microenvironment is remodelled during subclonal expansion, we next leveraged Slide-tags data to dissect the expansion-associated cell state transitions at single-nucleus resolution. After quantifying phylogenetic fitness on trees inferred from Slide-tags data, we stratified nuclei into high- and low-fitness groups and inspected the cell type abundances in their spatial neighbourhoods (**Fig. 3f**; **Extended Data Fig. 7a**; **Methods**). As expected, we found that the EMT cancer cell state was most consistently enriched in neighbourhoods surrounding high-fitness nuclei (**Fig. 3f**). With respect to differential enrichment of specific immune and stromal populations, we found that *Arg1+* TAM and myCAF populations were consistently enriched in spatial neighbourhoods of high-fitness cells whereas iCAFs and other TAMs were not (**Fig. 3f**).

Prompted by differential enrichment of TAM and CAF populations, we turned to systematically probing the cell state changes associated with subclonal expansions. Focusing on TAM and CAF populations, we performed differential expression of these cell types based on their spatial proximity to expanding cancer cells (**Extended Data Fig. 7b-c**). In addition to high *Arg1* expression, we found that TAMs in expanding tumour regions were characterized by the presence of the hypoxia-induced factor *Egnl3*, the immunosuppressive marker *Spp1*^67^, and programs indicating endocytosis and complement activity (**Extended Data Fig. 7b; Supplementary Table 3**). CAFs in expanding neighbourhoods were characterised by increased expression of genes implicated in hypoxia sensing, collagen synthesis, and fibrosis (**Extended Data Fig. 7c; Supplementary Table 3**). Likewise, in a more comprehensive analysis of the 44 Slide-seq arrays, we found that spatial neighbourhoods surrounding high-fitness, low-plasticity spots were most enriched for EMT, Hypoxic, and Fibrotic communities (C3, C10, and C5, respectively) and depleted for Alveolar, Endothelial, and Inflammatory communities (C1, C7, and C9, respectively) (**Extended Data Fig. 7d-e**; **Methods**).

These data suggest a model by which subclonal expansions towards a hypoxic tumour interior that, in turn, polarises non-tumour cells into pro-tumour, immunosuppressive states. Indeed, in returning to our previous Slide-seq analysis of community program assignments across tumour progression, we observed that the Hypoxic community (C10) appears prior to EMT (C3) when ranked by the transcriptional fitness signature (**Fig. 2f; Extended Data Fig. 5c**). In further support of this, immunofluorescence staining of KP-Tracer tumours revealed that hypoxia (as evidenced by the canonical hypoxia marker GLUT1 [*Slc2a1]* protein levels^71,72^) preceded the emergence of immunosuppressive ARG1+ immune cells (**Fig. 3g**). Importantly, we also found these same patterns in large surgical resections of human lung adenocarcinoma (**Methods**). Using immunofluorescence, we found significant co-localisation of hypoxia signalling and immunosuppressive SPP1+ protein levels (*p < 1e-5,* Fisher’s Exact Test); we also find that SPP1 and macrophage-marker CD68 overlap suggests that the majority of SPP1+ cells are macrophages (**Extended Data Fig. 7f; Methods**). These hypoxic, immunosuppressive tumour regions were also enriched with increased VIM expression, a marker of cancer cell EMT (*p < 1e-5,* Fisher’s Exact Test; **Extended Data Fig. 7g; Methods**). Consistent with these data, a reanalysis of published spatial transcriptomics data of human NSCLC tumours^40^ revealed that expression of the hypoxia-reporter *SLC2A1* (also known as *GLUT1*) was spatially associated with cell proliferation (as measured by *MKI67*), *TGF* 𝛽 signalling, EMT (*SNAI2*), and immunosuppressive macrophage polarisation (*SPP1, C1QB,* and *FCGR2B)* (**Extended Data Fig. 7h-i**).

### Differential intercellular interactions and hypoxia in the expanding niche drive cancer cell state changes

Our analyses thus far have revealed that expansion is characterized by a hypoxic, immunosuppressive niche that is accompanied by marked changes in cell type abundance and gene expression states. To gain additional molecular insight into these changes, we first investigated how cell-cell interactions differed between the expanding and non-expanding tumour niches. Applying a computational approach that identifies spatially localized cell-cell interactions to all our Slide-tags data, we found substantial reprogramming of interactions in expanding niches (**Fig. 3h; Extended Data Fig. 8**; **Supplementary Table 4; Methods**). This systematic analysis provided insight into the multitude of intercellular signalling pathways that integrate to establish and maintain the expanding niche: for example, from cancer cells, we find signals to macrophages that promote their recruitment (e.g., *Mif::Cd74*, *Mif::Cd44,* and *Il34::Csf1r*) and polarization to immunosuppressive states (e.g., *Bmp7*::*Acvr1*, *Mertk::Pros1*, and *Angptl2::Pirb*); in addition we find cancer cell-derived signals to fibroblasts that promote extracellular matrix remodelling and fibrosis (e.g., *Sdc1::Tnc, Fgf1::Fgfr1, Pdgfa::Pdgfrb*, and *Spp1::Itga8*). Reciprocally, cancer cells receive signals from this remodelled environment that promote the transition to proliferative and invasive phenotypes: in addition to pervasive TGF𝛽 signaling^73^ (a well-known inducer of EMT), cancer cells also received cell-type-specific signals from TAMs (e.g., *Fn1::Cd44*, *Il18::Il18r1,* and *Jag2::Notch2*) and CAFs (including several collagen and laminin genes signalling through *Cd44* and integrins on cancer cells). As orthogonal evidence of these interactions, we find that several are co-contained within the communities identified from the 44 Slide-seq arrays (e.g., *Fn1*, *Cd44*, *Spp1*, *Itga5* and *Bmp7* are all found in the EMT module, C3; **Supplementary Table 2; Extended Data Fig. 5a**). Together, this integrated spatial-lineage analysis provides a comprehensive catalogue of the multitudinous, and complementary, intercellular signals that synergize to establish a remodelled microenvironmental niche that promotes and stabilizes aggressive cancer cell states.

These differential intercellular signals in the hypoxic expanding niche led us to ask how various extrinsic signals drive cancer cell state changes and tumour progression. Because hypoxia has been shown to induce cell-state changes (including to EMT^74–76^), we tested the relative contributions of intercellular interactions with lung resident cells and hypoxia on major cancer cell states reported in this model^50,54,55^ (including AT2-like, gastric-like, high-plasticity, and EMT states). To model the complexity of the tumour microenvironment, we utilized a procedure to derive primary organoids from fresh KP tumours^77^ and co-cultured them in normal and hypoxic conditions with primary lung mesenchymal cells and alveolar macrophages isolated from wild type mice (**Fig. 3i**). In cancer cells, hypoxia alone induced a mild decrease in alveolar cell markers (*Sftpc* and *Cebpa*) and gastric-like markers (*Gkn2* and *Hnf4a*), a slight increase of the early mesenchymal marker (*Vim*), and no change in the key EMT transcription factor *Twist1*; on the other hand, co-culture of cancer cells with alveolar macrophages and lung mesenchymal cells under normal oxygen condition enhanced the epithelial identity of cancer cells (marked by upregulation of *Epcam* and lineage markers *Sftpc* and *Cebpa*) and the high-plasticity cell state marker *Slc4a11*^55^. Remarkably, when co-culturing cancer organoids, lung mesenchymal cells, and alveolar macrophages under hypoxia we observed a robust induction of EMT in cancer cells, with more than 2.5-fold increase of classical EMT markers (*Vim* and *Twist1*) compared to what is expected from hypoxia alone (**Fig. 3j; Supplementary Table 5**). Together, these data provide critical insights into the synergistic role between hypoxia and differential intercellular interactions in driving cancer cell state changes, highlighting that cancer cells must receive both external stimuli to enter and maintain an aggressive EMT-like state.

### Spatially resolved lineages reveal the evolution of metastasis-initiating niches in the primary tumour

Metastasis, the ultimate stage of tumour progression, accounts for approximately 90% of cancer-related mortality and is associated with pervasive microenvironmental remodelling^78–82^. However, it has remained challenging to delineate the specific microenvironmental features associated with tumour evolutionary dynamics during metastasis progression. Outstanding questions include: do the niches surrounding subclones giving rise to metastases differ from those surrounding other subclones? How do these gene expression programs change during metastatic spread? Our spatial-lineage platform is well-suited to identify the spatial localisation of metastasis-initiating subclones and characterize the microenvironmental remodelling associated with each step of the metastatic cascade.

We began by performing spatial transcriptomics on a KP-Tracer mouse with multiple primary lung tumours and widespread metastases in the mediastinal lymph node, rib cage, and diaphragm (**Fig. 4a, Extended Data Fig. 9a**). To maximise the probability of detecting metastasis-initiating subclones in primary tumours, we sampled multiple representative layers of the tumour-bearing lung at approximately 200-500um intervals, enabling us to study multiple large primary tumours from top-to-bottom and better track the clonal dynamics beyond a two-dimensional analysis. Tumour segmentation of Slide-seq data from these sections and coarse-grained spatial alignment determined by shared lineage states revealed four major tumours that could be tracked across layers (**Fig. 4b**).

**Figure 4.**
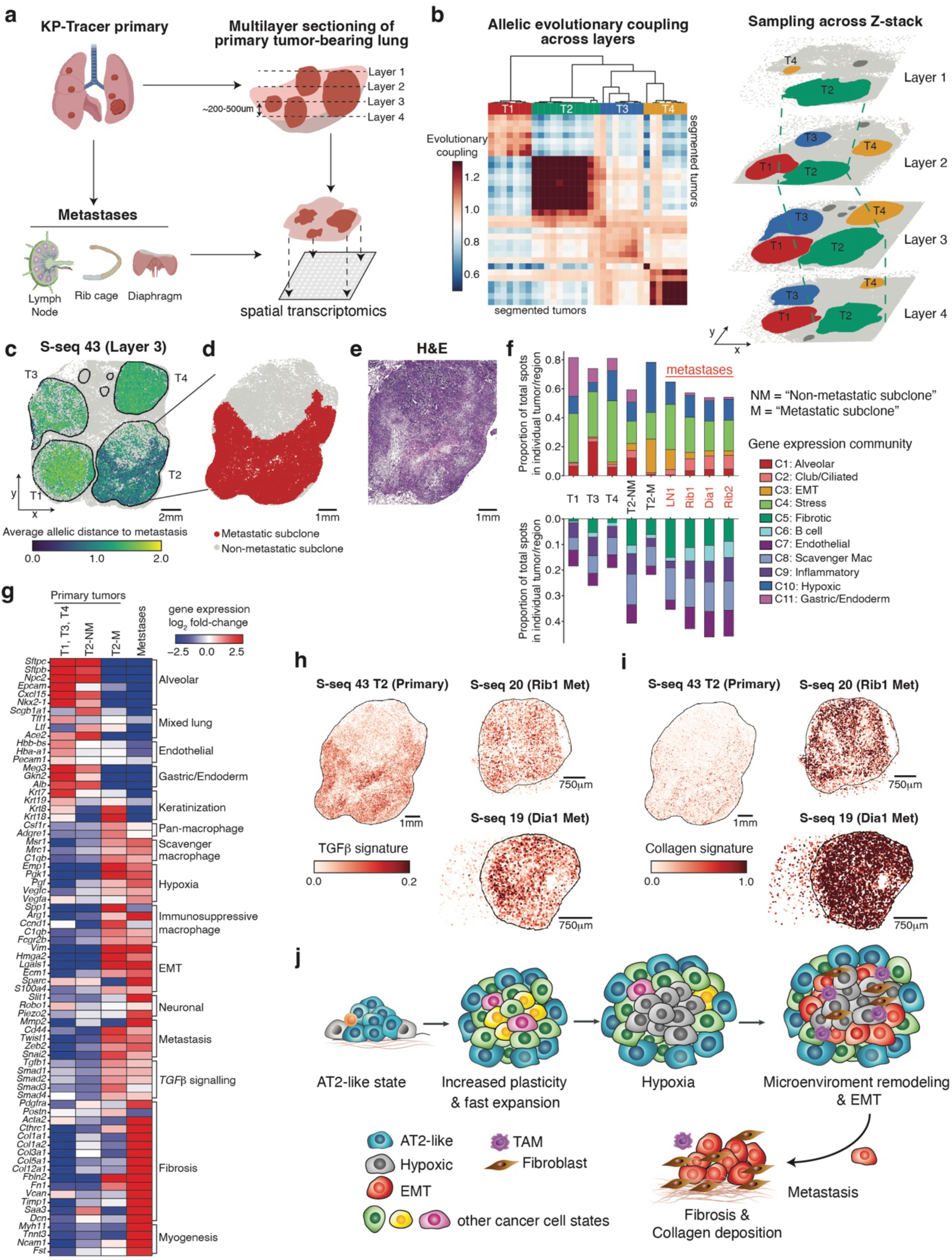
Tracing the evolution of subclonal niches across the metastatic cascade. **(a)** Schematic of spatial transcriptomics workflow from a KP-Tracer mouse with large primary lung tumours and paired metastases from the lymph node, rib cage, and diaphragm. Multiple lung sections with four large primary tumours were harvested and subjected to both Slide-seq and Slide-tags assays. Biorender was used to create parts of this schematic. **(b)** Coarse-grained alignment of Slide-seq spatial transcriptomics data (based on lineage-tracing edits) from four representative layers (Layer 1 – Layer 4) of a KP tumour bearing lung at approximately 200-500𝜇m intervals from different z position. (Left) A clustered heatmap of allelic evolutionary coupling scores across all Slide-seq datasets from the tumour-bearing lung identifies the four major tumours. Each row or column is a single tumour from one Slide-seq dataset. (Right) 3D reconstruction of aligned datasets, annotated by one of four major tumours. Individual tumours are labelled in different colours. **(c)** Representative spatial projection (S-seq 43) of allelic distances – summarizing how different lineage-tracing edits are between cells – for each spot with lineage-tracing data. Distance was computed to a consensus metastatic parental allele and normalised between 0 and 2. **(d-e)** The metastasis-initiating subclone in T2 was segmented from cells with high relatedness to metastatic tumours and labelled in red. **(e)** H&E staining of T2. **(f)** Proportion of gene expression community across representative stages of the metastatic cascade, including primary lung tumours (T1,3,4) without relatedness to metastases, the metastasis-initiating (M) and non-metastatic-initiating (NM) subclones in the primary tumour (T2) that gave rise to metastases, and four metastases. Top: communities that are more related to tumour or epithelial programs. Bottom: communities that are related to stromal/immune programs. **(g)** Heatmap of gene expression log2-fold-changes between environmental niche (primary tumours without metastatic relationship, non-metastasis-initiating (NM) and metastasis-initiating (M) subclones within T2, and metastases). Genes are manually organized into ontologies. **(h-j)** Spatial projection of gene expression scores of the Hallmark TGF𝛽 and custom Collagen gene signatures on the metastasis-initiating primary tumour and selected metastases. Tumour 2 on S-seq 43 is used as the representative layer. **(j)** A schematic model of KP tumour evolution and microenvironmental remodelling.

Our spatial-lineages in the large Slide-seq assays provide an opportunity to both compare the trajectory of multiple tumours and understand the transcriptional evolution of the niche surrounding the metastasis-initiating subclone in a single primary tumour. To do so, we first identified the spatial localisation of subclones giving rise to metastasis by inspecting the allelic similarities between primary tumours and metastases (**Fig. 4c; Extended Data Fig. 9b-c**). This analysis revealed that metastases from all 3 locations were phylogenetically related to a spatially coherent subclone in primary Tumour 2 (“T2”). Though distinct indels support this relationship between T2 and its related metastases (**Extended Data Fig. 9d**), the sparsity of spatial data and low lineage resolution within the expansion precluded an accurate estimation of the number of metastatic seeding events; however, because all metastases shared a common indel state, and metastasis is rare in this model^50,65^, a parsimonious explanation is that metastasis was initiated via a monoclonal seeding from T2 and subsequent secondary seeding events to different metastatic sites. Nevertheless, T2 could be identified in each layer independently and could be thus tracked across all sampled layers of this primary tumour (**Extended Data Fig. 9b-c**). This pattern was consistent in matched Slide-tags data (**Extended Data Figure 9c**), overlapped with subclonal expansions identified from our phylogenies, and was associated with regions exhibiting poorly differentiated histological features (**Fig. 4c-e; Extended Data Fig. 9e-f**). Leveraging the three-dimensional (3D) sampling of this tissue, and because all metastases shared indels with an expanding subclone that could be found across layers, we infer that it is likely that all metastases arose after subclonal expansion.

To understand the phylogenetic and gene expression programs underlying metastatic potential in this region of T2, we segmented this tumour into a niche surrounding the cells giving rise to metastases (“T2-Met”) or otherwise (“T2-NonMet”) and compared their gene expression patterns (**Fig. 4c-e**; **Methods**). The T2-Met niche had higher proportions of the EMT and Hypoxic communities (C3 & C10, respectively) and lower proportions of the Gastric/Endoderm and Alveolar communities (C11 & C1, respectively) (**Fig. 4f**). The T2-Met niche additionally down-regulated genes associated with Gastric and Endoderm states (e.g., *Gkn2* and *Meg3*), and had higher expression of genes marking cancer cell EMT (e.g., *Vim*), scavenger macrophages (e.g., *Mrc1* and *Msr1*), immunosuppressive macrophages (e.g., *Arg1* and *Fcgr2b*), TGF𝛽 signalling (e.g., *Tgfb1* and *Smad4*), and fibrosis (e.g., *Cthrc1* and *Postn*) (**Fig. 4g**). Orthogonal analysis with Slide-tags data corroborated these findings, as *Arg1+* TAMs and myCAFs were most enriched in spatial neighbourhoods of cells in the primary tumour related to metastases (**Extended Data Fig. 9g**). Moreover, immunofluorescence staining confirmed that ARG1+ cells co-localized with the metastasis-initiating VIM+ region of the T2 primary tumour (**Extended Data Fig. 9h**). Together, these results nominate several key molecular processes as potential drivers of the pro-metastatic niche, including fibrosis, TGF𝛽 signalling, and intercellular interactions between cancer cells, activated fibroblasts, and *Arg1+* immunosuppressive macrophages.

### Metastatic colonisation is accompanied by sustained EMT and enhanced fibrosis

Beyond the evolution within the primary tumours, we next investigated whether the microenvironments at distant metastatic sites are remodelled to resemble, or diverge from, the metastasis-initiating niche within the primary tumour. Comparing the niches surrounding metastases and the T2-Met subclone in the primary tumour, we found that metastases contained proportionally more regions annotated by stromal or immune communities and showed specifically higher representation of the Fibrotic community (C5) (**Fig. 4f**). As these communities represent several gene programs and may mask fine-scaled cell type changes, we further characterised the differential gene expression changes distinguishing niches of the primary tumour and metastases (**Fig. 4g**). While metastases up-regulated genes also found to distinguish the T2-Met niche – such as the EMT markers *Vim* and *Hmga2* and TGF𝛽-related genes – metastases displayed up-regulation of genes associated with collagen deposition (e.g., *Col1a1* and *Col12a1*) and myogenesis (*Tnnt3* and *Ncam1*) (**Fig. 4g**). After quantifying the activity of these gene expression programs in Slide-seq spots, we confirmed that these aggregated gene expression signals were spatially localised to tumour regions: metastatic tumours generally resembled the metastasis-initiating subclone in the primary tumour (for example with respect to TGF𝛽 signalling: log2FC = -0.14, t-test *p=*1.0; **Fig. 4h**) but substantially up-regulated collagen-related genes as compared to the primary tumour (log2FC = 3.81, t-test *p<1e-5*) (**Fig. 4i**). In a re-analysis of previously published datasets of metastatic KP tumours^50,65^, we also found that if EMT and TGF𝛽 signaling was present in the metastasis-initiating primary tumour, these programs were maintained (or enhanced) during metastasis across a range of distal sites (**Extended Data Fig. 10a**). Immunofluorescence staining further corroborated a marked increase in COL3A1 protein in metastases as compared to primary tumours **(Extended Data Fig. 10b**). Using our Slide-seq data, we found that TGF𝛽 signalling was significantly higher in metastases compared to surrounding normal tissue; however, collagen deposition was increased in half of our samples, indicating that metastases may co-opt the structure of the distal site to promote colonization (**Extended Data Fig. 10c**).

Extending our results beyond our mouse model, we additionally analysed single-cell RNA-seq data from a cohort of 56 human lung adenocarcinoma patients^83^ (including 34 brain metastases; **Extended Data Fig. 10d**). Our analysis revealed that distant metastases maintained high activity of the key gene expression programs we observed in our mouse data, including collagen deposition, TGF 𝛽 signalling, hypoxia sensing, immunosuppressive myeloid cell recruitment, and EMT (**Extended Data Fig. 10e-h**). Collectively, these results complement recent findings that TGF𝛽 signalling is critical for EMT and metastatic seeding^79^, and highlight that key properties of the metastasis-initiating tumour (e.g, EMT and fibrosis) are consistently preserved in remodelling of the distant site.

## DISCUSSION

In this study, we integrated high-resolution spatial transcriptomics with Cas9-based lineage tracing in a genetically engineered mouse model of lung adenocarcinoma to dissect the dynamic interplay between tumour evolution and microenvironmental remodelling in a spatially resolved fashion. Our analysis uncovered spatial communities associated with different stages of tumour progression; revealed relationships between tumour growth, plasticity and microenvironmental remodelling; and identified metastasis-initiating subclones that informed on the spatiotemporal evolution of gene expression along the metastatic cascade. These results present an unprecedented spatial map of lung adenocarcinoma evolution and the temporal ordering of key tumour evolutionary events, showcasing the power of integrating spatially resolved transcriptomics and single-cell lineage tracing to dissect the complex tumour dynamics underlying cancer progression.

This spatial-lineage platform provides new insights into our understanding of lung adenocarcinoma evolution in the KP model by revealing how key processes – such as subclonal expansion, hypoxia, and EMT – emerge and interrelate over time and space in the native microenvironment (**Fig. 4j)**. Our previous results provided several lines of evidence that tumours, following the initial loss of an AT2-like state, are characterised by a cancer-cell-intrinsic increase in clonal plasticity, leading to gains in transcriptional heterogeneity and subsequent subclonal expansion^50^. In the present study, we find that rapid subclonal expansion pushes early-stage cancer cells to the tumour periphery and contributes to the formation of a hypoxic microenvironmental niche. This hypoxic niche promotes additional microenvironmental remodelling characterised by *Arg1+* immunosuppressive myeloid subsets and myCAF-like fibroblasts; for example, by recruiting myeloid cells through hypoxia-induced chemokine secretion (e.g., *Ccl2, Ccl6,* and *Csf1*) and polarising immune and stromal cells through hypoxia-induced signalling cascades (e.g., *Hif1a* and *Vegfa*) as previously suggested^84–88^ (**Extended Data Fig. 4a; Extended Data Fig. 5a; Extended Data Fig. 7b-c; Supplementary Table 2-3**). In turn, our organoid experiments suggest that these reprogrammed immunosuppressive and pro-fibrotic cells integrate with hypoxia to drive cancer cell state transitions, leading to the emergence of a stable and pro-metastatic EMT state, for example through *TGF*𝛽 signalling as shown in our analysis (**Fig. 3h-j**; **Fig. 4g-h; Extended Data Fig. 9h**) and detailed in a recent study^79^. Upon dissemination, metastatic cells retain a mesenchymal phenotype (e.g., through sustained TGFβ signalling) and can further remodel the microenvironment into an enhanced fibrotic niche marked by increased collagen deposition (**Extended Data Fig. 10**).

Our proposed model of tumour progression in the KP lung cancer model has key implications into the temporal ordering of how cancer-intrinsic alterations and external signals integrate to regulate cancer cell states. Specifically, in the first stage of tumour progression, cancer cells enter a permissive epigenetic state characterized by increased plasticity and transcriptional heterogeneity but maintain their epithelial identity^50,54,55^. Our analysis of cell-cell interactions and 3D organoid co-culture experiments suggest that this epigenetic state is stabilized by normal, tissue-resident cells in the microenvironment (**Fig. 3h-k**; **Extended Data Fig. 8**). As high-plasticity regions of these tumours do not appear to be spatially coherent (**Fig. 3a**; **Extended Data Fig. 6i-k**), we propose that this first stage of tumour progression is mostly driven by cell-intrinsic epigenetic changes accompanying the loss of the AT2-like state.

In contrast, we find that after subclonal expansion, the tumour is substantially remodelled towards a hypoxic state characterized by immunosuppressive and fibrotic communities. Hypoxia has been shown to play critical roles in lung adenocarcinoma^89^ and other cancers (e.g., glioma^90^ and clear cell renal cell carcinoma^22^) where it is associated with genomic instability and EMT^22,74–76,91^. However, our understanding of how and when hypoxic conditions arise, and how hypoxia coordinates both further tumour microenvironmental remodelling and cancer cell state changes, have remained limited. Our model provides new insights into these questions, as we provide several lines of evidence that subclonal expansion actively contributes to the formation of a hypoxic niche within the tumour, which in turn drives the polarisation of immunosuppressive and fibrotic cell types that trigger distinct cancer cell state changes and induce the EMT state (**Fig. 2f**; **Extended Data Fig. 5c; Fig. 3e-j**). To note, our proposed model that multiple external stimuli (here, hypoxia and differential intercellular interactions from the remodelled tumour microenvironment) cooperate with cell-intrinsic changes in plasticity aligns well with data from other models, such as pancreatic adenocarcinoma^92^. Moreover, consistent with work demonstrating that EMT and metastasis are initiated in the perinecrotic tumour core^93^, we find that metastatic initiation occurs from spatially-confined subclones that have undergone EMT around the hypoxic niche (**Fig. 3a**; **Fig 4c-f**). With supporting evidence from our analysis of lung adenocarcinoma patient cohorts (**Extended Data Fig. 7f-h; Extended Data Fig. 10d-h**), we believe our model provides fundamental insights into the drivers and consequences of hypoxia, and more generally how feedback between intrinsic and extrinsic forces shape cancer cell states.

In addition to elucidating new aspects of how tumour evolution unfolds spatially, our study also sets the foundation for further studies. First, mechanistic studies will be needed to dissect how the hypoxic niche polarises immune and stromal subsets, and how this might lead to an aggressive, mesenchymal tumour state. As we have previously reported that plasticity plays an important role in tumour progression^28,50,55,94^, one area of research will be how hypoxia affects the high-plasticity cell states in lung cancer. Second, the platform we developed here can be adapted to study the spatiotemporal dynamics of tumour evolution in other models or under different perturbations. Notably, our platform is also amenable to modelling the effect of additional genetic perturbations as Cas9 is continuously expressed for tracing^50^. Third, while we introduced new computational approaches for phylogenetic reconstruction approaches that address the sparsity, resolution, and scale of these data, there remain opportunities to build new algorithms specifically tailored to the spatial aspect of data and statistically infer how spatial organisation affects phylogenetic patterns.

While our study reveals new aspects of tumour progression, there are limitations in the interpretation and extensibility of the approaches applied here. First, a single slide section may not represent the entirety of clonal dynamics in a tumour. To minimize this potential bias, we corroborated phylogenetic patterns with histology, orthogonal gene expression signatures derived from our previous single-cell lineage-tracing data (derived from unbiased sampling of whole tumours) and analysing representative sections at different depths of tumours from a tumour-bearing lung in **Fig. 4**. As scaling spatial transcriptomics experiments becomes more affordable, future studies can more densely sample three-dimensional structure to entirely account for this bias. Second, as a consequence of profiling tumour sections and the sparsity of spatial transcriptomics assays, we observe less indel diversity in spatial lineage tracing data than in previous applications with unbiased sampling, leading to lower resolution phylogenetic relationships. This may be ameliorated by optimizing the lineage-tracing kinetics and adapting tools for recording past molecular signalling events^95,96^. Third, the molecular sparsity and resolution of Slide-seq data pose a challenge in reconstructing phylogenies and detecting smaller spatial neighbourhoods (**Extended Data Fig. 2**). While we provide a spatial imputation algorithm to account for these technical issues, and benchmark its effectiveness in a variety of experiments from simulated and held-out real data, we anticipate that this imputation approach may have limitations in cases where lineage data is not spatially coherent, for example in systems with higher degrees of cell migration. In these scenarios, either alternative technologies with improved capture and resolution or new algorithms for performing spatial imputation and detecting robust spatial communities will be necessary. Finally, the trees presented in this study are only estimates of true phylogenetic relationships and may not truly reflect cell division histories; moreover, our quantifications of phylodynamic properties – expansion and plasticity – may be masked by non-uniform death rates and unobserved intermediate states. When possible, our study uses orthogonal data and approaches to substantiate all claims.

In summary, our study unites the insights provided by spatially resolved lineages and transcriptomics to investigate the fundamental patterns of tumour growth and its interactions with the microenvironment. Our analyses lead to a comprehensive model of how a tumour grows from a single, transformed cell into a large and complex ecosystem and provided new evidence for how tumour expansion-associated microenvironmental remodelling contributes to a distinct wave of cell state reprogramming towards pro-metastatic states. As one of the most comprehensive datasets of spatial tumour evolution to date, we anticipate that this resource will help pioneer new computational methods and quantitative and predictive models of tumour evolution.

## ACKNOWLEDGEMENTS

We thank Jack Rose, Can Ergen, Chen Weng, Pu Zheng, Sean-Luc Shanahan, Yun Zhang, Meaghan McGery, Santiago Naranjo, Theodore Esantsi, Raymond Ho, Michelle Chan, Romain Lopez, Adam Gayoso, and all members of the Weissman, Yang, Yosef, Chen, and Chang labs for helpful discussions. We thank Cristen Muresan, Anne Odera, Maria Gould, Daniel Braslavsky, Maxim Litvinov, and Nicole Dow for administrative support. We thank the Whitehead Institute and Broad Institute Sequencing Facility for sequencing support.

M.G.J. is supported by an NCI Pathway to Independence Award (NIH K99CA286968). N.M.A. was supported by a NIH F30 fellowship (1F30CA278495). K.E.Y. was supported by the National Cancer Institute of the National Institutes of Health under Award Number K00CA253729. L.W.K. is supported by a Helen Hay Whitney Postdoctoral Fellowship. T.J. laboratory currently also receives funding from The Lustgarten Foundation for Pancreatic Cancer Research, but this funding did not support the research described in this manuscript. This work was supported in part by the Cancer Center Support (core) grant P30-CA14051 from the National Cancer Institute and by the NIH grant R35CA274464. T.J. is the David H. Koch Professor of Biology and a Daniel K. Ludwig Scholar. J.S.W. is supported by the Howard Hughes Medical Institute, NCI Cancer Target Discovery and Development (CTD^2) and NIH Centers of Excellence in Genomic Science (CEGS). Both J.S.W. and T.J. received fundings from Ludwig Center at MIT. D.Y. is supported by a Damon Runyon Dale Frey Award, an NCI Transition Career Development Award 1K22CA289207, an NIH Director’s New Innovator Award 1DP2OD037078, and a Lung Cancer Research Foundation Leading Edge Award. N.Y. is supported in part by an NIH grant R56-HG013117 and by the European Union Council (ERC, Tx-phylogeography, 101089213). Views and opinions expressed are however those of the authors only and do not necessarily reflect those of the European Union or the European Research Council Executive Agency. Neither the European Union nor the granting authority can be held responsible for them. F.C. acknowledges support from NIH Early Independence Award (DP5, 1DP5OD024583), the NHGRI (R01, R01HG010647), the Burroughs Wellcome Fund CASI award, the Searle Scholars Foundation, the Harvard Stem Cell Institute, and the Merkin Institute. This research was supported by the NYSCF. FC is a New York Stem Cell Foundation – Robertson Investigator. B.I. was supported by the National Institutes of Health (NIH) through National Cancer Institute (NCI) grants R01CA266446, R01CA280414, R37CA258829, and U54CA274506; B.I. was additionally supported by a Velocity Fellows Award, the Louis V. Gerstner, Jr. Scholars Program, a Tara Miller Team Science Award Metastasis Research by the Melanoma Research Alliance, the Pershing Square Sohn Cancer Research Alliance Award, and the Herbert Irving Comprehensive Cancer Center (HICCC) Human Tissue Immunology and Immunotherapy Initiative. B.I. is a Cancer Research Institute Lloyd J. Old STAR (CRI5579). E.D. was supported by an NIH F30 award (1F30CA301910-01).

## AUTHOR CONTRIBUTIONS

D.Y., J.S.W., N.Y., F.C. and K.H.(J.)M. conceived of the project. D.Y., L.T, and D.S. transduced mice, sacrificed mice, harvested tumours, and constructed spatial transcriptomics sequencing libraries. W.M.R III generated the KP-Tracer chimeric mice. M.G.J. analysed the lineage-tracing and gene expression spatial transcriptomics data with help from K.H.(J.)M., W.N.C, and V.Z.C. M.G.J. and K.H.(J.) M. performed simulation benchmarks for Slide-seq data with input from W.N.C., L.W.K., and K.E.Y. J.W. performed spatial-mapping of Slide-tags data. D.S. and N.M.A. performed staining and imaging of H&E and immunofluorescence and histology analysis. T.T and M.D. developed the ligand-receptor detection algorithm and T.T. performed ligand-receptor analysis with help from M.G.J. and D.S. H.W., and E.C performed organoid coculture experiments and gene expression analysis with supervision from D.Y. and J.L. E.D. and B.I. provided access to PBMA data and performed analysis on LUAD brain metastases. B.H, and A.S. collected and identified human LUAD surgical resections. G.M. and V.K. aided with synthesizing pucks for spatial transcriptomics experiments. D.Y., M.G.J., D.S., T.J., J.S.W., N.Y., and F.C. interpreted the results. M.G.J., D.Y., and D.S. wrote the manuscript with input from all authors.

## DATA AND CODE AVAILABILITY

Custom code and tutorials for the analysis of spatially-resolved lineage-tracing data is available on Github through Cassiopeia (https://github.com/YosefLab/Cassiopeia). Notebooks and code for reproducing analyses are available at https://github.com/mattjones315/KPSpatial-release. All raw and processed data will be made available on GEO and other public repositories.

## SUPPLEMENTARY TABLES

**Supplementary Table 1**: Meta data for samples

**Supplementary Table 2**: Genes in spatial communities.

**Supplementary Table 3**: Fitness-neighbourhood differential expression and GO Term analyses.

**Supplementary Table 4**: LARIS ranked cell-cell communication list.

**Supplementary Table 5**: Quantitative PCR measurements for organoid experiments.

## METHODS

### EXPERIMENTAL MODELS AND SUBJECT DETAILS

#### KP-Tracer mice

KP-Tracer mouse was generated by generating chimeric mice from blastocyst injection of engineered, lineage tracer enabled mouse embryonic stem cells harbouring conditional alleles *Kras^LSL-G12D/+^;Trp53^fl/fl^*; *Rosa26^LSL-Cas9-P2A-mNeonGreen^* as previously described^50^. Eight-to-twelve-week-old KP-Tracer mice were infected intratracheally with ad5-SPC-Cre virus (1x10^8 Pfu) purchased from University of Iowa viral vector core for tumour initiation. This enables specific tumour initiation and lineage-tracing in Alveolar Type II (AT2) cells, the major cell-type of origin of lung adenocarcinoma. All studies were performed under an animal protocol approved by the Massachusetts Institute of Technology (MIT) and Columbia University Committee on Animal Care. Mice were assessed for morbidity according to MIT and Columbia Division of Comparative Medicine guidelines and humanely sacrificed prior to natural expiration.

#### Surgical resection of human lung adenocarcinoma

Formalin fixed paraffine embedded tissue specimens of human lung adenocarcinoma surgical resections were acquired from the Columbia University Molecular Pathology Core Facility. These samples were initially collected under Institutional Review Board (IRB)-approved protocol AAAA3987 at New York Presbyterian Hospital/Columbia University Medical Center (CUMC), and all processing procedures performed on patient samples were in accordance with the ethical standards of the IRB. Twelve surgical resections of KRAS-mutant lung adenocarcinoma were used for immunostaining analysis.

### METHODS DETAILS

#### Sample processing

Tumour-bearing lungs were harvested and re-inflated with ∼2ml of 50% OCT (1:1 mix with PBS) and 1:100 of RNase inhibitor (NEB M0314L). After cleaning up excess blood and liquid, the whole tissue was embedded in 100% OCT and frozen using dry ice-methanol bath. Frozen samples were kept at -80C until sectioning for further analysis.

#### Spatial transcriptomics with Slide-seqV2

##### For 3 mm and 5.5 mm arrays

Fresh frozen tissues were cryo-sectioned at a thickness of 10 μm using a Cryostat (CM1950, Leica) set at −17 to −18 °C. The tissue sections were carefully transferred onto precooled arrays, which were placed on top of a glass slide inside the cryostat. A finger was briefly placed underneath the slide to melt the tissue and adhere it to the array. Immediately after, the tissue and array were transferred together into a 1.5 ml or 2 ml Eppendorf tube containing 200 μl (for 3 mm arrays) or 500 μl (for 5.5 mm arrays) of hybridisation buffer (6x SSC with 2 U μl^−1^ Lucigen NxGen RNase inhibitor, Lucigen, 30281). The samples were incubated in the hybridisation buffer for 15 minutes to 1 hour at room temperature, allowing the RNA to bind to the oligonucleotides on the beads. After incubation, the tissue and array were briefly dipped into 1x Maxima RT buffer to wash off the hybridisation buffer and then transferred to the reverse transcription (RT) reaction mixture (1x Maxima RT buffer, 1 mM dNTPs (NEB, N0477L), 2 U μl^−1^ Lucigen NxGen RNase inhibitor, 2.5 μM template switch oligonucleotide, 10 U/μL Maxima H Minus reverse transcriptase (Thermofisher Scientific, EP0753)). The tissue and array were incubated in 200 μl (for 3 mm arrays) and 500 μl (for 5.5 mm arrays) of the RT reaction mixture for 30 minutes at room temperature, followed by 1.5 hours at 52 °C. To digest the tissue, 200 μl (for 3 mm arrays) or 500 μl (for 5.5 mm arrays) of tissue digestion buffer (100 mM Tris-Cl pH 7.5, 200 mM NaCl, 2% (w/v) SDS, 5 mM EDTA and 16 U ml^−1^ proteinase K (NEB, P8107S)) was added to the reaction mixture and incubated at 37 °C for 30 minutes. Following digestion, 200 μl (for 3 mm arrays) or 500 μl (for 5.5 mm arrays) of wash buffer (10 mM Tris pH 8.0, 1 mM EDTA and 0.01% Tween-20) was added, and a P200 pipette was used to carefully triturate the beads off the array. The beads were centrifuged at 3000g for 2 minutes, followed by three washes with wash buffer. To remove RNA strands, the beads were incubated in 0.1N NaOH for 5 minutes, followed by a wash with wash buffer and 1x TE buffer, and centrifuged again at 3000g for 2 minutes. Second-strand synthesis was performed by mixing the beads with 200 μl (for 3 mm arrays) or 500 μl (for 5.5 mm arrays) of second-strand synthesis mixture (1x Maxima RT buffer, 1 mM dNTPs, 10 μM dN-SMRT oligonucleotide and 125U ml^−1^ Klenow enzyme (NEB, M0210L)) and incubating at 37 °C for 1 hour. The beads were then washed three times with wash buffer and once with water. cDNA amplification was carried out by resuspending the beads in 200 μl (for 3mm arrays) or 1.2 ml (for 5.5 mm arrays) of cDNA amplification mixture (1x Terra Direct PCR mix buffer (Takara Biosciences, 639270), 25U ml^-1^ of Terra polymerase (Takara Biosciences, 639270), 2 μM TruSeq PCR handle primer and 2 μM SMART PCR primer). The reaction was divided into 50 μl aliquots and amplified using the following PCR program: 95 °C for 3 min; four cycles of 98 °C for 20 s, 65 °C for 45 s and 72 °C for 3 min; nine cycles of 98 °C for 20 s, 67 °C for 20 s and 72 °C for 3 min; 72 °C for 5 min; hold at 4 °C. The cDNA product was purified twice using SPRI beads (Beckman Coulter, B23318) at a 0.8x bead-to-sample ratio, eluting in a final volume of 20 μl (for 3mm arrays) and 60 μl (for 5.5 mm arrays). A total of 1 ng (for 3 mm arrays) or 3x 1ng (for 5.5 mm arrays) of cDNA was used for Illumina sequencing library construction. The Nextera XT kit (Illumina, FC-131-1096) was used for tagmentation, followed by amplification with TruSeq5 and N700 series barcoded index primers. Libraries were cleaned with SPRI beads according to the manufacturer’s instructions at a 0.6x bead-to-sample ratio and resuspended in 10 μl of water per reaction. Lineage tracing target site libraries were amplified from cDNA and prepared fpr Illumina sequencing using previously described protocols^50^.

##### For Curio 1 cm arrays

The buffers and enzymes used were the same as those described for the 3 mm and 5.5 mm arrays but adjusted for scale. In brief, hybridisation, dipping, washing, RT reaction and tissue digestion were performed using the reservoirs provided by Curio with 500 μl volume for each step. After tissue digestion the beads were divided into 2 tubes for wash buffer washes and combined for cDNA amplification. A total of 4.8 ml of cDNA amplification mixture was prepared, and the reaction was divided into 50 μl aliquots for cDNA amplification in 96-well PCR plates, following the same PCR program as outlined previously. cDNA was purified twice using 0.8x SPRI beads and eluted in a final volume of 80 μl. 8x 1ng cDNA products were used for Illumina sequencing library preparation through tagmentation with a Nextera XT kit, followed by amplification and cleanup as stated above. Lineage tracing target site libraries were amplified from cDNA and prepared fpr Illumina sequencing using previously described protocols^50^.

#### Spatial transcriptomics with Slide-tags

Fresh frozen tissues were cryo-sectioned at 20 μm thickness using a Cryostat set at −17 to −18 °C. Precooled 6 mm square custom-made biopsy punches were used to punch and isolate regions of interest from the tissue sections. The isolated tissue regions were carefully transferred onto a precooled array, which was placed on top of a glass slide. A finger was briefly placed underneath the slide to melt the tissue onto the array. Immediately after, the tissue, array, and slide were placed on ice, and approximately 10 μl of dissociation buffer (82 mM Na_2_SO_4_, 30 mM K_2_SO_4_, 10 mM glucose, 10 mM HEPES, 5 mM MgCl_2_) was gently pipetted onto the tissue to ensure it was fully covered. The array was then exposed to an ultraviolet (UV) light source (0.42 mW mm^-2^, Thorlabs, M365LP1-C5, Thorlabs, LEDD1B) for 1 minute to cleave spatial barcode oligonucleotides off the beads. After photo-cleavage, the array was incubated on ice for 7.5 minutes before being transferred to a well of a 12-well plate. To release the tissue from the array, 1 ml of extraction buffer (dissociation buffer with 1% Kollidon VA64, 0.2% Triton X-100, 1% BSA, 666 U ml^-1^ RNase-inhibitor) was gently dispensed onto the array, and the buffer was carefully triturated up and down over the tissue 10–15 times. This process was repeated until the tissue was completely released from the array. The array was then discarded, and mechanical dissociation of the tissue was performed by triturating the supernatant 100–150 times using a 1 ml pipette to fully release the nuclei from the tissue. The extraction buffer containing the nuclei was transferred to a 15 ml tube. The well was washed three times with 1 ml of wash buffer (dissociation buffer with 1%BSA and 1: 100 RNase-inhibitor) and the washes were pooled into the same 15 ml tube. The final volume of the wash buffer was adjusted to 10 ml. The nuclei were centrifuged at 600g for 10 minutes at 4 °C. After centrifugation, 9.5 ml of the supernatant was carefully removed. The pellet was resuspended and passed through a precooled 40 μm cell strainer (Corning, 431750) into a 1.5 eppendorf tube. The 15 ml tube and cell strainer were washed with 1 ml of wash buffer, and the nuclei were pelleted again by centrifuging at 600g for 10 minutes at 4 °C. After centrifugation, the supernatant was carefully removed, leaving approximately 50 μl of wash buffer for nuclei resuspension. To determine cell count, 2 μl of resuspended nuclei was mixed with 18 μl of 1: 100 diluted DAPI, and the nuclei were manually counted using a C-Chip Fuchs-Rosenthal disposable hemocytometer (INCYTO, DHC-F01-5). Based on the cell count, up to 25,000 nuclei were processed using the Chromium Next GEM Single Cell 3’ Reagent Kits v3.1 (with Feature Barcode technology for Cell Surface Protein, 10x Genomics, PN-1000268). Lineage tracing target site libraries were amplified from cDNA and prepared fpr Illumina sequencing using previously described protocols^50^.

#### H&E staining

H&E was performed with a Leica ST5010 Autostainer XL and Leica CV5030 Fully Automated Glass Coverslipper. Bright-field images were taken using the Leica Aperio VERSA Brightfield, Fluorescence & FISH Digital Pathology Scanner under a ×10 objective. Tumour grade was analysed in H&E-stained sections using an automated deep neural network developed by Aiforia.

#### Sequencing

Sequencing was performed at using NovaSeq S4. For Slide-seq gene expression libraries: read1: 50bp, read2: 50bp, index1: 8bp was used. For Slide-seq Target Site libraries: read1: 44bp, read2: 260bp, index1: 8bp was used. For Slide-tags gene expression libraries: read1: 28bp, read2: 90bp, index1: 10bp, index2: 10bp was used. For Slide-tags gene expression libraries: read1: 28bp, read2: 260bp, index1: 8bp setting was used.

#### Immunofluorescence staining & imaging

15 μm-20 μm tissue sections were fixed in 4% PFA at room temperature for 10-15 min. The sections were washed twice in 1x PBS. Antigen retrieval was performed by boiling 1X IHC Antigen Retrieval Solution (ThermoFisher Scientific, 00-4955-58) and incubating tissue sections inside for 30 min until the solution cooled down, followed by washing tissue sections with 1x PBS and incubated in 0.3% PBST (0.3% Triton X-100 in PBS) at room temperature for 10 min. Three times of 1x PBS wash was then performed. Blocking (0.5% BSA and 0.1% Triton X-100 in 1x PBS) was performed at room temperature for 1 hour. Tissue sections were incubated with primary antibodies: VIM (1: 200, Biotechne, AF2105), CD31 (1: 200, Biotechne, AF3628), ARG1 (1: 200; Cell Signaling Technology, 93668), GLUT1 (1: 100; AbCam, ab195020), CD45 (1: 200, Cell Signaling Technology, 70257), and COL3A1 (1: 200, Proteintech, 22734-1-AP) at 4 °C overnight. Tissue sections were washed three times with 1x PBS and further incubated with secondary antibodies (donkey anti-goat 405, 1: 1000, ThermoFisher Scientific, A-48259; donkey anti-mouse 647, 1: 1000, ThermoFisher Scientific, A-31571; donkey anti-rabbit 647, 1: 1000, ThermoFisher Scientific) at room temperature for 2-3 hours. Tissue sections were then washed three times with 1x PBS, mounted and imaged using Dragonfly 201-40 High Speed Confocal Imaging Platform.

#### Mouse lung tissue dissociation for isolation of mesenchyme and alveolar macrophages

For isolation of mesenchymal cells and alveolar macrophages, mice were culled by cervical dislocation and lungs perfused with 10ml of cold PBS. Mouse lungs were inflated via intratracheal injection with 3ml of dispase solution (Fisher Scientific, 11553550), containing Collagenase I (GIBCO, 17100017) at 350U/ml. Lungs were carefully dissected out of the thoracic cavity, placed in a 50ml falcon and minced into small pieces. Cells were washes down with 3ml of PBS and samples were places in a shaking incubator at 37°C, 190 rpm, for 45 min. 7.5ul of 1% DNase I (Sigma, D4527) was added to each sample in the final 10 min of incubation. The resulting cell suspensions were filtered through 100 µm and 40 µm cell strainers and washed with 2 ml of 10% fetal bovine serum (FBS) in PBS (PF10). Samples were centrifuged at 800 rpm for 5 min at 4°C and pellets were resuspended in 1 ml of red blood cell lysis buffer (RBC buffer, Merck, 11814389001) for 60 seconds at RT. After lysis, 6 ml of Dulbecco’s Modified Eagle Medium Nutrient Mixture F-12 820 (DMEM/F-12, Invitrogen, 11330057) was added to neutralize the RBC buffer. 500µL of filtered FBS was added slowly to the bottom of each tube and samples were centrifuged again at 800rpm for 5 min at 4°C. Cell pellets were resuspended in PF10 and placed into Eppendorf tubes for antibody staining.

#### Primary 3D mouse lung organoid co-cultures

14 weeks post tumor initiation by Ad-SPC-Cre, tumour cells from the *Kras^LSL-G12D/+^; Trp53^fl/fl^; R26^LSL-Cas9-P2A-GFP^*mice were dissociated and isolated (CD45^-^ CD31^-^ GFP^+^) via fluorescence-activated cell sorting (FACS) as previously described^50^. Mesenchymal cells (CD45^-^ CD31^-^ EpCAM^-^) and alveolar macrophages (CD45^+^ CD64^+^ SiglecF^+^) were isolated from adult wild type C57BL/6 mice. The following antibodies were used: APC Anti-Mouse CD45 (559864, BD Biosciences); PE/Cyanine7 anti-mouse CD64 (139313, Biolegend); PE anti-mouse CD170 (Siglec-F) (155505, Biolegend); APC Anti-Mouse CD31 (551262, BD Biosciences); PE/Cyanine7 anti-mouse CD326 (Ep-CAM) (118216, Biolegend).

Using sorted cells, tumour organoid co-cultures were stablished according to previous publications78. Briefly, freshly sorted tumour and stromal cells were centrifuged at 300g for 10 min at 4°C and resuspended in 3D basic media composed by DMEM/F12 (GIBCO, 11330-032) supplemented with 10% of FBS and Insulin-Transferrin-Selenium (ITS, Corning, 25-800-CR). Cells were counted and combined to obtain mixtures of 5k tumour cells with 45k mesenchyme and 30k alveolar macrophages per well. Cells were centrifuged at 300g for 10 min at 4°C, resuspended in 100 μl GFR-Matrigel (Corning, 356231) containing 50% 3D basic medium and platted in a 24-well Transwell insert with 0.4 μm pore (Greiner, 662641). 500 μL of 3D basic medium was added in the lower chamber and cells were cultured at 37°C with 5% CO2. Cultures containing 5k of tumour cells were also stablished, following the same protocol. 10 µM Rho kinase (ROCK) inhibitor Y-27632 (Cayman Chemical. #10005583) was added to all cultures for the first 48h. After the initial 48h, cells were transferred to either 37°C with standard 20% oxygen (Normal oxygen) or to 37°C with 5% oxygen (Hypoxia) and were maintained for an additional 9 days, with media changed every other day. At the end of the experiment, cultures were dissociated and GFP+ cells sorted and stored for RNA extraction. Total RNA was isolated from the sorted cancer cells (∼20k cells per sample) using PicoPure RNA isolation Kit (ThermoFisher Scientific, KIT0204). cDNA was generated using High-Capacity cDNA Reverse Transcription Kit (ThermoFisher Scientific, 4368814). Quantitative PCR was performed using PowerTrack SYBR Green Master Mix (ThermoFisher Scientific, A46012). All qPCR primers are listed in **Supplementary Table 5**.

#### Human LUAD FFPE sample staining and quantification

Formalin-fixed, paraffin-embedded (FFPE) human lung adenocarcinoma (LUAD) tissue sections were first incubated at 70 °C for 1 hour. Deparaffinization was then performed using a Leica ST5010 Autostainer XL with the following settings: 60 °C incubation for 15 min, followed by 3x xylene washes for 3 min each, 2x 100% ethanol washes for 3 min each, 1x 95% ethanol wash for 3 min, and water wash for 5 min. Sections were subsequently subjected to antigen retrieval by de-crosslinking in a vegetable steamer for 30 min using Antigen Retrieval Buffer (pH 6.0, Abcam, ab93684) and were then cooled to room temperature. Permeabilization was performed by incubating the sections in 0.3% PBST (0.3% Triton X-100 in 1x PBS) for 10 min at room temperature, followed by three washes in 1x PBS. Blocking was carried out for 1 hour at room temperature using a blocking buffer containing 0.5% BSA and 0.1% Triton X-100 in 1x PBS. Sections were incubated overnight at 4 °C with primary antibodies. For quantifying GLUT1, SPP1, and CD68 signals, anti-GLUT1-Alexa Fluor 647 (1:100; Abcam, ab195020) and anti-CD68-Alexa Fluor 555 (1:200; CST, 23308) were used; anti-SPP1 (1:200; Merck, HPA027541) was applied to adjacent sections. For quantifying GLUT1 and VIM signals, anti-GLUT1 (1:100; Abcam, ab195020), anti-VIM (1:200; Biotechne, AF2105), and anti-ACTA2 (1:250; Thermo Fisher Scientific, MA1-06110) were used. For unconjugated primary antibodies (e.g., SPP1), sections were washed three times in 1x PBS and incubated for 2–3 hours at room temperature with appropriate secondary antibodies: donkey anti-rabbit Alexa Fluor 594 (1:1000; Thermo Fisher Scientific, A-21207), donkey anti-goat Alexa Fluor 405 (1:1000; A48259), donkey anti-rabbit Alexa Fluor 647 (1:1000; A32795), and donkey anti-mouse Alexa Fluor 488 (1:1000; A32766). Nuclei were counterstained with DAPI or Propidium Iodide (PI), followed by three 1x PBS washes. Finally, sections were mounted and imaged using a Dragonfly 201-40 High-Speed Confocal Imaging Platform.

For quantification, 3-5 randomly selected 2 x 2 tiled scan images at 10x magnification were captured from annotated tumor regions, as determined by prior H&E staining and pathologist review. Each 2 x 2 tiled image was divided into four quadrants: top-left, top-right, bottom-left, and bottom-right. For SPP1 and GLUT1 quantification, these subregions were assessed for GLUT1 and SPP1 signal positivity, and regions were classified into four categories: GLUT1^+^SPP1^+^, GLUT1^+^SPP1^-^, GLUT1^-^SPP1^+^, and GLUT1^-^SPP1^-^. For quantifying VIM signal associated with tumors, only VIM^+^/ACTA2^-^ regions were considered VIM-positive, while VIM^+^/ACTA2^+^ regions were classified as VIM-negative.

### QUANTIFICATION AND STATISTICAL ANALYSIS

#### Slide-seqV2 gene expression quantification and quality-control

A python implementation of Kallisto-bustools^97^ (*kb_python,* version 0.27.3 available at https://github.com/pachterlab/kb_python) was used for transcript quantification and processing from raw FASTQs produced with Slide-seq. Specifically, we utilised the *count* procedure implemented in Kallisto that quantifies the number of UMIs in a Slide-seq library that map to each transcript sequence in the provided reference (here, *mm10*). To account for the unique read structure of the Slide-seq library, we invoked the *count* procedure with the flag -x “0,0,8,0,26,32:0,32,41:1,0,0”. To determine a whitelist of barcodes to use during quantification, we matched barcodes identified with kallisto to the spatial barcodes and their coordinates observed during *in situ* sequencing of the Slide-seq array during fabrication^56,57^. We then used a custom script to assign spatial coordinates, identified during *in situ* sequencing of the Slide-seq array prior to running the assay, to quantifications from the kallisto pipeline and returned an AnnData structure containing the spatially-resolved transcript abundances for each spot. To supplement the barcode filtering during the kallisto pipeline, we applied an extra filter requiring at least 150 UMIs observed in a spot. For most analyses, we utilise log-normalised counts where each cell’s UMI total is scaled to the median library size and a log1p transformation is applied. When scaled counts are used, we additionally use Scanpy’s *scale* function with a max value of 10.

#### Slide-tags gene expression quantification and quality-control

Similar to Slide-seq processing, we utilised the python implementation of Kallisto-bustools^97^ (*kb_python,* version 0.27.3 available at https://github.com/pachterlab/kb_python) to quantify transcript abundance from FASTQ data. As this data represents reads from sequencing single-nuclei with the 10X V3 kit, we utilised the *--umi-gene*, *--workflow nucleus*, and *-x 10XV3* flags. Similar to the Slide-seq analysis, we utilised the mm10 transcriptome reference.

After transcript quantification, we applied several quality-control procedures. First, we removed background gene expression signal from ambient RNA by applying Cellbender^98^ (version 0.3.0, available at https://github.com/broadinstitute/CellBender) to the unfiltered gene expression counts. We used default settings for all libraries, except for 10X Library 9 where we used the following flags: --empty-drop-training-fraction 0.15, --total-droplets-included 20000, -- learning-rate 0.0001, and --epochs 300. After running Cellbender, we applied further cell-filters to remove outliers with high mitochondrial or ribosomal content (between 5-15% for libraries). We further inspected the count distribution in each library and removed nuclei with excessively high UMI content (approximately 20,000 UMIs). All quality-control was performed with Scanpy^99^ (version 1.10.0, downloaded via *pip*). For most analyses, we utilise log-normalised counts where each cell’s UMI total is scaled to the median library size and a log1p transformation is applied. When scaled counts are used, we additionally use Scanpy’s *scale* function with a max value of 10.

#### Slide-seq lineage tracing target-site data processing

To begin processing target-site data, we trimmed reads from Slide-seq libraries using cutadapt^100^ (version 4.1) with the following flags: -m :250 --max-n 0.2 --discard-untrimmed -O 10 --no-indels --match-read-wildcards -e 2 -j 16 --action retain -G AATCCAGCTAGCTGTGCAGC. We then applied Cassiopeia^41^ (version 2.0.0, available at https://github.com/YosefLab/Cassiopeia) to trimmed FASTQs using the “slideseq2” chemistry and specific parameters for Slide-seq libraries. First, to account for the possibility of multiple cells observed in a given spot, we allowed allele conflicts (*allow_allele_conflicts = True*) and did not enable doublet filtering. While we performed intBC whitelist correction, we did not perform additional error correction to remove intBCs with conflicting alleles (this is similarly motived by the fact more than one cell can be observed in a given spot). We additionally relaxed the UMI/cell threshold to account for reduced capture of Slide-seq assays (*min_umi_per_cell = 2*). Finally, we utilised the “likelihood” method for UMI collapsing, with *max_hq_mismatches =* 3 and *max_indels =* 2. Other settings remained default. This pipeline produced a cleaned allele table, reporting the set of intBCs and alleles for each observed spot, that was used for tree reconstruction.

#### Slide-tags lineage tracing target-site data processing

Cassiopeia^41^ (version 2.0.0, available at https://github.com/YosefLab/Cassiopeia) was used to process FASTQs containing target-site data. As Slide-tags represents single-nucleus data, we utilised default settings except for a more relaxed UMI/cell cutoff (*min_umi_per_cell = 5)* to reflect the reduced sensitivity of single-nucleus sequencing. As a part of default settings, we corrected cell barcodes to those observed after quality-control filtering, corrected intBCs to a whitelist for the corresponding mESC (E1) with a distance threshold of 1, and performed UMI (with a maximum distance of 2) and intBC error correction (minimum UMI support of 5) to correct for conflicting target sites observed in the same nuclei. Doublets were filtered out using the default conflicting threshold of 35%. This pipeline produced a cleaned allele table, reporting the set of intBCs and alleles for each observed spot, that was used for tree reconstruction.

#### Slide-tags spatial barcode processing

Spatial mapping of Slide-tags nuclei was achieved as previously described^58^. Briefly, reads from spatial barcode FASTQ files were filtered for those containing the spatial barcode universal primer constant sequence and cell barcode sequences from a called cell barcode whitelist generated by the gene expression pipeline (see above section entitled “Slide-tags gene expression quantification and quality-control”). Spatial barcode sequences were matched with a whitelist of in situ sequenced spatial barcodes, assigning spatial coordinates to each true spatial barcode. The set of spatial barcodes and the corresponding x,y coordinates for each cell barcode were clustered with DBSCAN^101^ (implemented in the R package *dbscan*, version 1.1−11). For cell barcodes with a single cluster of spatial barcodes, spatial barcodes not contained in the cluster were filtered out and a UMI-weighted centroid of the remaining spatial barcodes represented the x,y coordinates of the cell barcode. DBSCAN parameters were determined from a sweep of minPts values (3 to 15) under a constant eps = 50. The chosen minPts positioned the highest proportion of cell barcodes.

#### Spatial imputation of lineage-tracing data

To recover lineage-tracing data for reconstruction on spatial assays, we performed spatially-informed imputation of target site data. To begin, we first created a character matrix from the allele tables constructed from target-site lineage tracing processing. In this character matrix, denoted as Χ, each row corresponds to a cell (or spot) and each column corresponds to a particular cut site in an integration barcode (intBC). For clarity of notation, we refer to each cut-site/intBC pair as a character, and thus in our system a character matrix will have (|intBCs| x 3) columns. The entry Χ[𝑖, 𝑗] denotes the edit (which we refer to as a “state”) observed at the *i*^th^ cell/spot in the *j^th^* character. The missing data rate refers to the proportion of entries in this character matrix that do not have data that pass our quality-control filters.

To perform spatial imputation, we first constructed a spatial nearest-neighbour graph (𝑁) such that each spot was connected to all other spots within 30𝜇m of the spot. For each missing entry in character matrix, 𝑖, 𝑗 we queried the frequency of states at character j in all neighbours of spot i in 𝑁. If the concordance of a particular state was higher than 80% in these neighbours, then we replaced the entry 𝑋[𝑖, 𝑗] with this state. To minimize the effect of nearby stromal cells in a neighbourhood – which should not have active lineage-tracing – we did not allow this state to be 0, the uncut state. To maximise the alleles were used during spatial imputation, we required each state to be supported by at least 3 UMIs. We reported this procedure for each missing entry in the character matrix for a total of 5 iterations which continued to remove missing data from the character matrices with no apparent reduction in accuracy in simulations or held-out real data (**Extended Data Fig. 1j-n**).

#### Benchmarks of imputation and reconstruction accuracy

To benchmark the accuracy of spatial imputation and downstream effects on tree reconstruction, we utilised two different strategies:

- <u>Synthetic data</u>: First, we utilised the Cas9-based lineage-tracing simulation framework in Cassiopeia^41^ (version 2.0.0, available at https://github.com/YosefLab/Cassiopeia). Specifically, we simulated trees using Cassiopeia’s *BirthDeathSimulator* with the following parameters: 5000 extant cells, and utilised a LogNormal birth-waiting distribution parameterised by log (𝑓) where *f* is a fitness coefficient that accumulates with each cell division (in each cell division, a new coefficient 𝑓 ∼ 𝑁(0, 0.25) is drawn and added to the base fitness) and a standard deviation of 0.5. Then, we simulated lineage tracing data onto the tree with Cassiopeia’s *Cas9LineageTracingDataSimulator* with desired mutation proportion of 0.7, 100 states, 39 cut sites (representing our system with approximately 13 intBCs, each with 3 cut-sites), and no missing data rates at this point. Then, we simulated spatial coordinates on each tree using the *ClonalSpatialDataSimulator* over a shape of (1,1,1) and sampled a 2D slice from this 3D simulation at random. Finally, we subsampled from this spatial array using the *UniformLeafSubsampler* in Cassiopeia with a rate of 0.4 (resulting in lineages with 2,000 observations) and induced random dropout at various rates: [0.1, 0.25, 0.5, 0.6, 0.7, 0.9]. We simulated 10 trees for each parameter combination. As the spatial array simulated does not exactly match that from Slide-seq, we applied a modified *k-nearest-neighbour* graph construction approach, linking together spots to their closest 10 neighbours and performed spatial imputation (see section titled “Spatial imputation of lineage-tracing data”). We required concordance of 0.8 for the selected state and at least 5 votes. Since this simulated data does not include any normal cells, we do allow the imputation of the state 0. We reported the accuracy of this imputation strategy in **Extended Data Fig. 2e**). Then, we compared the tree reconstructing accuracies using the *triplets_correct* function in Cassiopeia for reconstructions with or without imputation and for different reconstruction strategies: modified Neighbour-Joining, Cassiopeia-Greedy, or a hybrid of these two approaches (see section “Phylogenetic reconstruction”).
- <u>Simulated held-out Slide-seq data:</u> In the next experiment, we assessed the accuracy of recovering target-site data that was held-out from real Slide-seq data. To do this, for a given Slide-seq array, we masked out 10% of the observed data (supported by at least 3 UMIs) and performed spatial imputation in neighbourhoods of 30𝜇m using the strategy described previously (see section titled “Spatial imputation of lineage-tracing data). Similarly, we required a concordance of 0.8 and at least 5 votes in support of the imputed allele. We only considered samples where at least 10 states were imputed. Random predictions were obtained by shuffling the node labels in the neighbourhood graph. We reported the average accuracy and total number of imputed values over five replicates in **Extended Data Fig. 2h**.

#### Simulation benchmarks of lineage-tracing pre-processing

As a feature of the Slide-seq is that multiple cells may be observed in one spot^57^, multiple conflicting alleles can be observed for a given target site in a single spot. Typically, this would break the assumption of the Cassiopeia reconstruction pipeline (in single-cell approaches, we assume that only one allele can be tied to a given intBC and perform error correction or filtering otherwise). However, we implemented new reconstruction algorithms that can handle multiple conflicting states in each spot (see section entitled “Phylogenetic reconstruction”) and simulated the effects of various pre-processing techniques.

First, we simulated trees on two-dimensional surfaces where various proportions of cells would be grouped together based on their spatial location. To do so, we simulated simple binary trees of 2000 cells and overlaid lineage-tracing data with Cassiopeia’s *Cas9LineageTracingDataSimulator* function using the following parameters: 39 characters, a mutation proportion of 0.5, and no missing data. We then merged together cells using Cassiopeia’s *SupercellularSampler* method with rates of [0.1, 0.2, 0.3, 0.4, 0.5, 0.6]. We simulated 32 replicates.

For each replicate, we pre-processed character matrices according to three strategies. Here, the entry of the *i^th^* cell and *j^th^*character (denoted as Χ[𝑖, 𝑗]) would contain a set of states X[𝑖, 𝑗] = {𝑠_1_, 𝑠_2_, …, 𝑠*_k_*}, each state occurring at some frequency f(𝑠*_i_*) = 𝑓*_i_*. In the first strategy (“collapse duplicates”) we take the unique set of states so that X[𝑖, 𝑗] = {𝑠_1_, 𝑠_2_, …, 𝑠*_k_*′} 𝑠. t. 𝑓*_i_* = 1 ∀ i ∈ 𝑘′; in the second strategy (“most common”) we take the most common state, such that X[𝑖, 𝑗] = 𝑎𝑟𝑔𝑚𝑎𝑥*_f_*(*s*′)∀∈*k*𝑠′; and the third strategy (“all states”) we do not perform any filtering. In **Extended Data Fig. 2a** we report the tree reconstruction error (measured with normalised Robinson-Foulds distance) for trees reconstructed with Neighbour-Joining^63^.

#### Phylogenetic reconstruction on Slide-seq data

To enable phylogenetic reconstruction on Slide-seq data in which multiple cells can be contained in a single spot and thus conflicting alleles are present, we implemented a Hybrid Cassiopeia-Greedy & Neighbour-Joining algorithm that could utilise conflicting allele states.

For Cassiopeia-Greedy, we modified the splitting decision rule to account for all states observed in a spot. Cassiopeia-Greedy is a simple, heuristic-based algorithm for reconstructing phylogenies that iteratively finds the most common state in a given population and splits samples into groups based on the presence or absence of the state. It is based on a perfect-phylogeny reconstruction algorithm^102^ and has an efficient runtime of *O(mn)* for a population of *n* samples and *m* characters. Here, we changed the procedure to find the state with the highest frequency by allowing each sample to carry multiple states in a character. The runtime of this algorithm is still polynomial in the size of the sample population – *O(n(ms))* where in the worst case scenario every single state is observed in every single character; given the size of the spatial array, this is exceedingly uncommon and typically 1-3 cells are captured per spot^57^.

For Neighbour-Joining, we utilised the standard algorithm^63^ but with a modified distance map that accounts for multiple states per spot. Specifically, we implemented a new dissimilarity metric that takes in two sets of states 𝑆_1_ and 𝑆_2_ and computes all the pairwise allelic dissimilarities and reports a linkage similar to hierarchical clustering. Here, we use the modified allelic dissimilarity for two states 𝑠*_i_*, 𝑠*_j_* to compute distances between pairs of states, previously described^41,47,50^:

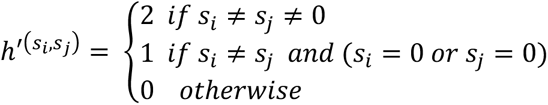

In the case where weights are passed in, then the dissimilarity function is computed as follows:

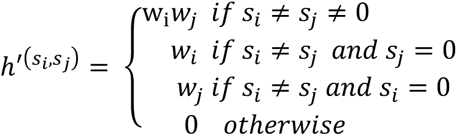

Then, we utilised a single linkage function such that only the smallest modified allelic dissimilarity across all pairs of states in 𝑆_1_ and 𝑆_2_ was used. This is to maintain such that if the same state is observed in two spots, the dissimilarity returned is 0.

For the hybrid reconstruction, we utilised the modified Cassiopeia-Greedy algorithm described above until subpopulations of size 1000 cells were found, at which point Neighbour-Joining with the modified dissimilarity metric was used to resolve phylogenetic relationships. We utilised state probabilities inferred from all Slide-seq and Slide-tags datasets and used the weight − log(𝑝*_i_*) for state 𝑠*_i_* during tree reconstruction.

#### Phylogenetic reconstruction on Slide-tags data

We utilised the standard Cassiopeia-Hybrid^41^ algorithm for reconstructing Slide-tags phylogenies. Briefly, this approach applies the heuristic-based Cassiopeia-Greedy algorithm to reconstruct relationships between the major subclones and then applies the maximum-parsimony-based Cassiopeia-ILP algorithm to solve fine-grained phylogenetic structure in smaller populations. As previously described in detail^41^, Cassiopeia-ILP proceeds by building a potential graph of all possible ancestral states (constrained in size by a user-defined parameter) and solves for the maximum-parsimony phylogeny by reconstructing a Steiner Tree on this data structure. The Steiner Tree problem is solved via an Integer Linear Program (ILP) allowed a certain time to converge. Here, the transition between Cassiopeia-Greedy and -ILP algorithms is determined by the distance to the latest common ancestor (LCA) of a subpopulation.

We applied the Cassiopeia-Hybrid algorithm with state priors inferred from all samples^41,47,50^, determined the switch between Greedy and ILP algorithms using an LCA cutoff of 20, devised a potential graph of 10000 nodes with a maximum distance of 15 across nodes (maximum_potential_graph_lca_distance=15), and allowed the ILP 12600s to converge.

#### CNV detection and analysis

inferCNV (https://github.com/broadinstitute/infercnv, version 1.18.1) was used to detect CNVs from Slide-tags or Slide-seq expression data. For Slide-tags data, inferCNV was used in “subcluster” mode with leiden resolution of 0.0005 and the following parameters: HMM=TRUE, BayesMaxPNormal=0.2, min_cells_per_gene = 1, cutoff = 0.1. Otherwise, default parameters were retained.

For Slide-seq data, owing to the sparsity in transcript capture, we performed inferCNV analysis on pooled gene expression data. Specifically, we combined the gene expression data for the adjacent spots into bins of size 50𝜇m (akin to 10X Visium data). Then, inferCNV was used in “subcluster” mode with leiden resolution of 0.0005 with the following parameters: window_length = 201, HMM=TRUE, BayesMaxPNormal=0.5, min_cells_per_gene = 1, and cutoff=0.2. Otherwise, default parameters were retained.

For counting the number of CNVs for the benchmarking analysis in **Extended Data Fig. 3a**, we used the HMM CNV predictions (*Pnorm_0.2.pred_cnv_regions.dat). To assess the Nearest Neighbour Purity between inferCNV data and lineage-tracing trees, we computed the fraction of leaves whose nearest neighbor shared the same inferCNV grouping (as reported in infercnv.observation_groupings.txt of the inferCNV pipeline). We removed inferCNV groupings with fewer than 20 cells. This purity score was compared to an empirical background in which inferCNV cluster labels were shuffled 1000 times.

#### Slide-tags cell type annotation

After performing quality-control on Slide-tags gene expression data, we assigned cell types first by integrating Slide-tags data with an annotated single-cell gene expression reference dataset of KP-Tracer tumours^50^ with scANVI^103^. To do so, first identified 4,750 variable genes using Scanpy’s^99^ *highly_variable_genes* function using the *flavor=“seurat_v3”* and raw counts. We then trained an scVI model^104,105^ on the joint dataset and these variable genes using 3 layers and 70 latent dimensions over 1000 epochs. Then, we transferred labels from the single-cell reference dataset to the Slide-tags nuclei with scANVI utilizing 200 samples per label and 100 epochs. Through this, we used the *gene_likelihood=“nb”* setting in training models and used the technology – Slide-tags or single-cell – variable to signify batch.

After training this model, subset to the scANVI embeddings to the Slide-tags data only and re-clustered the data with Scanpy^99^ using the Leiden algorithm^106^ and resolution 1.2. We then split clusters into those that appeared to derive from tumour/epithelial cells or those that derived from the stroma. To call tumour or epithelial clusters, we evaluated if a cluster had an abundance of tumour nuclei (defined as nuclei with target site data and at least 20% of their sites containing indels) or expressed the epithelial-lineage marker *Nxk2-1*. Immune cell clusters were identified based on the marker *Cd45 (Ptprc)* and other stromal cells were identified by expression of *Pdgfra, Col1a1,* or *Col5a1* (fibroblasts) or *Pecam1* (endothelial cells). For each subsetted dataset (tumour/epithelia or stromal), we reclustered the data and annotated cell types based on annotations predicted with scANVI and differentially expressed genes identified with Scanpy’s *rank_genes_group* function (using the Wilcoxon test).

#### Assessment of Slide-tags tumour cell type signatures in previous KP-Tracer data

To test the portability and accuracy of the tumour clusters identified in Slide-tags, we assessed the activity of gene signatures in the previous KP-Tracer data^50^. Specifically, we for each cell-type identified in Slide-tags, we computed the top 100 differentially-expressed genes using the Wilcoxon test in Scanpy^99^ and further filtered genes to have a log-fold change > 1 and an FDR-corrected p-value <= 0.01, and an AUROC of at least 0.6. We then used these genes to define a transcriptional signature for each Slide-tags cell type. each of these signatures, we scored the activity in cell types identified in Slide-tags data and the previous KP-Tracer dataset using the *score_genes* function using *n_bins=30* and *ctrl_size* equal to the number of genes in the gene set. Signatures were computed on scaled, log-normalised counts. The result of this analysis is presented in **Extended Data Fig. 4b**.

#### Slide-seq spatial community detection and scoring

To identify spatial communities in Slide-seq data, first applied the Hotspot^107^ algorithm for detecting spatially autocorrelated gene sets on each sample. In the spatial mode, this algorithm constructs a nearest neighbour graph based on spatial coordinates, computes an autocorrelation statistic for each gene, and then identifies modules of genes that have significant pairwise autocorrelation values. Here, we applied Hotspot with 20 neighbours, and FDR threshold of 0.01 to identify spatially autocorrelated genes, and a minimum module size of 50 genes.

Then, to identify robust modules of genes that appear across tumours, we assessed the Jaccard overlap between all pairs of modules across all tumours and filtered out modules that did not have a Jaccard overlap of at least 0.2 with at most one other module. We then performed Z-normalization on these Jaccard statistics and clustered these using hierarchical clustering (using the “ward” method on Euclidean distances) and identified 11 clusters, representing robust spatial modules.

As these robust modules are collections of modules across all samples we analysed, we distilled these down to a set of genes – representing what we call a “spatial community” in this study – by taking genes that appear in at least 25% of the modules in the robust module. Using these genes in the spatial community, we compute the activity of these communities for each spot (termed “community scores”) using the *score_genes* function in Scanpy^99^ with *ctrl_size=100* and *n_bins=30.* We computed these scores on scaled, log-normalised gene expression counts. To obtain community assignments for each spot, we took the community with the highest score.

#### Tumour segmentation

To segment tumours, we utilised the SpatialData^108^ package and the napari-spatialdata viewer for interactive annotation. To identify tumour areas on a sample, we overlaid phylogenetic subclones and the number of target-site UMIs detected and manually segmented areas that appeared to be (a) phylogenetically related and (b) had elevated target-site UMIs indicative of tumour regions. We saved these annotations and used the segmentations to perform downstream analysis on a tumour-by-tumour basis.

#### Fitness signature calculation

To quantify fitness signature scores, we utilised a gene set that was found to be associated with changes in fitness from our previous single-cell KP-Tracer study^50^. Using this gene set, we quantified the transcriptional activity for each spot in Slide-seq data by applying the *score_genes* function in Scanpy^99^ with *ctrl_size=100* and *n_bins=30.* We computed these scores on scaled, log-normalised gene expression counts.

#### Phylogenetic fitness inference

We quantified fitness on Slide-seq and Slide-tags phylogenies by utilizing the *LBIFitness* fitness estimator in Cassiopeia^41^. This function wraps a fitness estimator based on the “local branching index” as previously described^109^. This procedure has been previously used in our system^50^. Primed by the true single-cell resolution of Slide-tags trees, we estimated branch lengths using the *IIDExponentialMLE* branch length estimator in Cassiopeia. This function implements a function that provides maximum-likelihood branch lengths on a tree topology given the pattern of edits observed in the leaves and an assumptions about the irreversibility of Cas9 editing^110^. Using the branch lengths determined by this maximum-likelihood procedure, we estimated single-cell fitness on Slide-tags trees.

Due to the increased missingness on Slide-seq trees and the fact that MLE-based branch length approaches have not been benchmarked on Slide-seq data, we performed a more conservative branch length estimation, as done previously^50^. Here, branches had a length of 1 if they had any mutations along them, otherwise they had a branch length of 0. Using these branch lengths, we estimated single-cell fitness on Slide-seq trees.

#### Single-cell clonal plasticity quantification

To estimate single-cell clonal plasticity on phylogenies, we applied approaches described in our previous studies^50,68^. Specifically, on Slide-tags data where we have true single-cell data and associated cell type identities, we applied the *score_small_parsimony* procedure to all nodes in a tree using *meta_item=“cell_type”* and normalised by the number of leaves in the subtree induced by the node. Then, we computed plasticity for each cell by averaging together all the normalised parsimonies.

Since we do not have true single-cell resolution for Slide-seq data, we employed the L2 plasticity score described in our previous study^50^, using community scores. Specifically, let 𝐶*_i_* be the vector of community scores associated with spot 𝑖. For this spot 𝑖 we found its closest phylogenetic neighbours (denoted by set *N*) and then computed the L2-Plasticity (𝐿2*_p_*(𝑖)) for this spot by the average Euclidean distance to the vector of community scores for these neighbours:

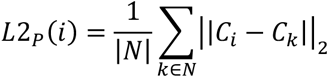

All scores were unit scaled.

#### Differential expression and abundance in neighbourhoods of high-fitness cells

To identify changes in gene expression and spatial communities associated with fitness, we first stratified cells into high- and low-fitness groups. In Slide-seq data, we computed single-cell fitness scores (see section above entitled “Phylogenetic fitness inference”) and identified a threshold separating two modes using *scipy.signal.argrelmin* in the merged fitness distributions and split spots into high-fitness groups and low-fitness groups based on this threshold. Only tumours with at least 200 observations with lineage-tracing data were used. As each fitness distribution is normalised within individual tumours to be unit-scaled, this approach finds a global pattern in high- and low-fitness cells. Then, we constructed a neighbourhood graph connecting each spot to all other spots within 30 𝜇 m. The community scores for all communities were computed in these neighbourhoods and the distributions in neighbourhoods of high- and low-fitness cell were reported in **Extended Data Fig. 7d**.

In Slide-tags data, high and low-fitness cells were similarly determined from the distribution of all fitnesses using *scipy.signal.argmin*. As Slide-tags is sparser than Slide-seq, we constructed neighbourhoods using the closest 20 cells (an example is shown in **Extended Data Fig. 7a**). We then identified the differentially-expressed genes in neighbourhoods of high- and low-fitness cells of all Macrophage and Fibroblast subsets using the t-test as implemented in Scanpy’s^99^ *rank_genes_groups* function. For the Macrophage analysis, we evaluated the Alveolar Macrophages, *Arg1+* TAMs, *Pecam1+* TAMs, and *Vegfa+* TAMs; for the Fibroblast analysis we evaluated the *Wt1+* fibroblast, iCAF-like and myCAF-like populations. Genes expressed in fewer than 50 cells were filtered out, and the differential expression statistics for the top 10,000 genes were computed. Genes with an absolute log2-fold-change > 1 and an FDR-corrected p-value < 0.01 were marked as significantly differentially expressed. To compute enrichments in these neighbourhoods, we computed the frequency of cell types in neighbourhoods of high- and low-fitness cells and divided by the expected fraction of these cell types given the overall distribution and size of the Slide-tags array.

GO Term analysis of differentially-expressed genes was performed using gseapy^111^ (version 1.1.3) with the following gene sets: “WikiPathways_2019_Mouse”, “Reactome_2022”, “GO_Biological_Process_2023”, “GO_Molecular_Function_2023”, and “KEGG_2019_Mouse”. Significant terms are reported in **Supplementary Table 3**.

#### Differential expression in neighbourhoods of high-plasticity cells in Slide-seq

Similar to the fitness-based analysis (see section entitled “Differential expression in neighbourhoods of high-fitness cells”), we stratified cells into high- and low-plasticity groups. After quantifying the L2-clonal plasticity score in Slide-seq data, we determined a threshold separating high- and low-plasticity regions if a cell had greater plasticity than the 60^th^ percentile or less than the 40^th^ percentile, respectively. Then, we constructed a neighbourhood graph connecting each spot to all other spots within 30𝜇m. The community scores for all communities were computed in these neighbourhoods and the distributions in neighbourhoods of high- and low-plasticity cells were reported in **Extended Data Fig. 7e.**

#### Cell-cell communication analysis of Slide-tags data

Ligand–receptor (LR) and cell type level interaction analysis was performed with Ligand And Receptor In Spatial Transcriptomics (*LARIS*) on the Slide-tags dataset. LARIS is a computational tool for spatial LR activity inference implemented in Python, capable of producing single cell, cell-type, or tissue region-level predictions in seconds. LARIS is based on the *in silico* spatial diffusion of ligands and receptors to simulate intercellular communication. Each single cell (or bead) with a known spatial location is assigned an LR score based on the expression of ligands and receptors in the nearest neighbours. This integrated LR score can be used for easier visualization of interactions in space. To capture spatial interaction patterns that show unique localization, cosine similarity is employed on the observed LR activity patterns against shuffled cell locations. This yields a ranked list of LR interactions that have localized spatial expression patterns, helping identify potentially interesting interactions cell type (or other) label-free. The cell type level prediction scores are calculated by combining the average of the LR integration scores with the specificity of ligands and receptors for each cell-type combination, resulting in a ranked list of cell type to cell type communication based on interaction strength and specificity. To identify region-specific interactions independently of cell type, regional labels (e.g., cancer clones) can be assigned to cells, and statistical tests or other approaches can be applied to the LR scores to identify differentially expressed ligand and receptor pairs.

Raw counts were first log-normalized using the Scanpy implementation of *normalize_total (target_sum=10000)* and *log1p* functions. The latest mouse *CellChatDB*^112^, formatted for *LARIS*, served as the reference for known LR pairs.

To calculate integrated LR scores, which quantify the predicted interaction strength based on receptor and ligand gene expression in spatially proximate cells, we applied the *calculateLigandReceptorIntegrationScore* function from *LARIS* (*number_nearest_neighbors=20*). Interactions exhibiting spatial specificity and variability compared to randomly shuffled locations were identified using the *runZoneTalk* function (*number_nearest_neighbors=20, mu=0.2, sigma=100*). To infer and rank interactions between each cell type combination, we employed the *calculateZoneTalkScoreByCellType*function (*mu=100, expressed_pct=0.05, mask_threshold=1e-8*), which considers the abundance, specificity, and spatial localization of each interaction. Visualisations of the -log10(interaction scores) of the top cell-type-specific interactions are displayed in **Extended Data Figure 8** and reported in **Supplementary Table 4**.

To detect differentially expressed LR pairs between expanding and non-expanding clones, each cell was labeled by both cell type and clonality. The COSG tool^113^, a cosine similarity–based marker gene identification method (*mu=1000, expressed_pct=0.15*), was applied to LR scores at both regional clonality and cell type levels. Visualization of the resulting LR scores was performed using Scanpy.

#### Coarse-grained alignment of Slide-seq data

To track the three-dimensional structure of clones across sampled layers in **Fig. 4**, we utilised the non-imputed processed target-site data (see section entitled “Slide-seq lineage tracing target-site data processing”). To maximise fidelity of slide registration, we enforced hard quality-control cutoffs, requiring each spot be supported by at least 7 UMIs and then subsequently each intBC-allele to be supported by at least 5 UMIs. We filtered out spots that had less than 20% of their sites reporting indels, or more than 70% missing data. We then computed modified allelic distances (see section above entitled “Phylogenetic reconstruction on Slide-seq data”) between all pairs of spots across layers. Modified allelic distances here are normalised by the number characters shared between two spots (thus are normalised to values between 0-2). For computational reasons, we did not allow ambiguous alleles (taking only the most frequent allele per intBC in a spot) as the distance calculation is memory- and time-intensive. Using this distance matrix, we computed allelic evolutionary couplings using *compute_evolutionary_coupling* function in Cassiopeia with the following parameters: *minimum_proportion = 0.0002, number_of_shuffles = 100*. We then normalised the evolutionary coupling as previously described^50^, as so:

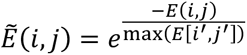

Where *E(i,j)* denotes the allelic evolutionary coupling between spot *i* and *j* and *max(E[i’, j’])* indicates the maximum value across all evolutionary couplings. Clusters identified via hierarchical clustering of the normalised allelic evolutionary coupling matrix were used as registered Tumour IDs in **Fig. 4b**.

#### Detection of metastasis-initiating subclones

To detect metastasis-initiating subclones in primary tumours, we created a shared character matrix between all lung sections profiled with 1cm x 1cm Curio arrays and Slide-seq samples of metastases. We filtered out spots that did not have at least 2 UMIs intBC-alleles that were not supported by at least 2 UMIs. We further filtered out spots that had fewer than 20% of their target-sites cut and more than 70% missingness. For computational reasons, we did not allow ambiguous alleles (taking only the most frequent allele per intBC in a spot) as the distance calculation is memory- and time-intensive. We then computed a shared metastatic parental allele state by taking states that were shared amongst 60% of spots in metastases profiled with Slide-seq. From this parental state, we computed the modified allelic distance (normalised by the number of shared characters) to all spots in the lung sample. We performed a similar analysis in paired Slide-tags data, computing the normalised modified allelic distances from all nuclei to the metastatic parental allele state.

#### Differential expression across metastatic cascade

We identified gene expression changes across niches associated with the metastatic cascade by employing the distances computed in the section above entitled “Detection of metastasis-initiating subclones”. We identified the metastasis-originating subclone as localizing to T2, so T1, T3 and T4 were determined to be Primary tumours without any relationship to the metastases. Focusing on T2, we further segmented it into a metastasis-initiating subclone (T2-Met) and other subclones (T2-NonMet). Specifically, we assigned cells to a metastatic subclone if their normalised modified allelic distance was less than 0.8. Then, using these assignments, we performed watershed segmentation with a custom procedure. Specifically, we binned signal into bins of 100 adjacent spots, applied a Gaussian filter with a sigma of 1.5 (with the Python package *skimage*) and then applied an Otsu threshold and dilation. We then applied an exact distance transform with *scipy.ndimage.distance_transform_edt* and computed a Watershed mask over peaks identified with *skimage.feature.peak_local_max* with a goal of identifying one tumour. This segmented subclone was labeled as T2-Met, and the remainder of the tumour was called T2-NonMet. We then performed differential expression across the library-size-normalised, logged counts of four groups (Primary tumours without metastatic relationship; T2-Met; T2-NonMet; and metastases) using a t-test implemented in Scanpy’s^99^ *rank_genes_groups* and reported the log2- fold-change in **Fig. 4g**.

Signature scores for TGF 𝛽 signalling were computed using MSigDB’s “HALLMARK_TGF_BETA_SIGNALING” signature^114^. Signature scores for collagen were computed for a custom gene set consisting of *Acta2, Col1a1, Col2a1, Col3a1, Col5a1,* and *Col12a1*. Significance was computed using a one-sided *t*-test assessing if signature scores were higher in the metastatic tumour as compared to the primary tumour.

#### Differential cell type abundance in metastatic neighbourhoods

Similar to analyses stratifying Slide-tags cells into neighbourhoods of high- and low-fitness cells, we stratified cells into neighbourhoods of cells closely related to metastases. As with determining cells related to metastases in Slide-seq data, we computed the distance to the parental metastatic allele and assigned cells with distances smaller than 0.8 as related to metastases. Then, we reconstructed spatial neighbourhoods of the closest 20 cells and quantified cell type enrichments based on the frequencies of cell types in these neighbourhoods and the overall frequency in a Slide-tags array.

#### Analysis of transcriptional signatures in metastatic tumours

To assess the transcriptional signatures of metastasis, we explored three additional datasets: (1) scRNA-seq from metastatic families (i.e., primary seeding tumour and related metastases) from our previous KP-Tracer study^50^; (2) bulk RNA-seq from several KP tumours at different stages, including early stage, non-seeding primary tumours, seeding primary tumours, and metastatic tumors^65^; and (3) single-cell RNA-seq from primary lung adenocarcinoma (LUAD) tumours and metastatic brain tumours from the Pan-cancer human Brain Metastases Atlas (PBMA)^83^.

In the re-analysis of KP-Tracer metastasis families^50^, we focused on the two families from the sgNT genotype (3724_NT_T1 and 3513_NT_T1). 3724_NT_T1 had 4 related metastases: 3 from the liver and 1 in soft tissue; 3513_NT_T1 had two related metastases: one in the kidney and one in lymph tissue. Data was pre-processed with Scanpy and we performed analysis on count-normalized (1M), logged, and scaled data. Signature gene sets were obtained from the Hallmark set in MSigDB^114^.

In the bulk analysis of KP tumors^65^, we obtained gene expression data from the original publication in Supplementary Table 1 and present gene expression for a selection of genes in **Extended Data Figure 10a**.

For the analysis of the PBMA study, we subset the data for an analysis of treatment-naïve LUAD tumours (n=22) and brain metastases (n=36; a total of 54 tumours). Data was pre-processed with Scanpy and we performed analysis on count-normalized (1M), logged, and scaled data. Signature scores for TGFb signalling and Hypoxia were computed using MSigDB’s Hallmark gene signature set^114^. Signature scores for collagen were computed for a custom gene set consisting of *ACTA2, COL1A1, COL1A2, COL3A1, COL5A1, COL12A1*.

## EXTENDED DATA FIGURES

**Extended Data Figure 1.**
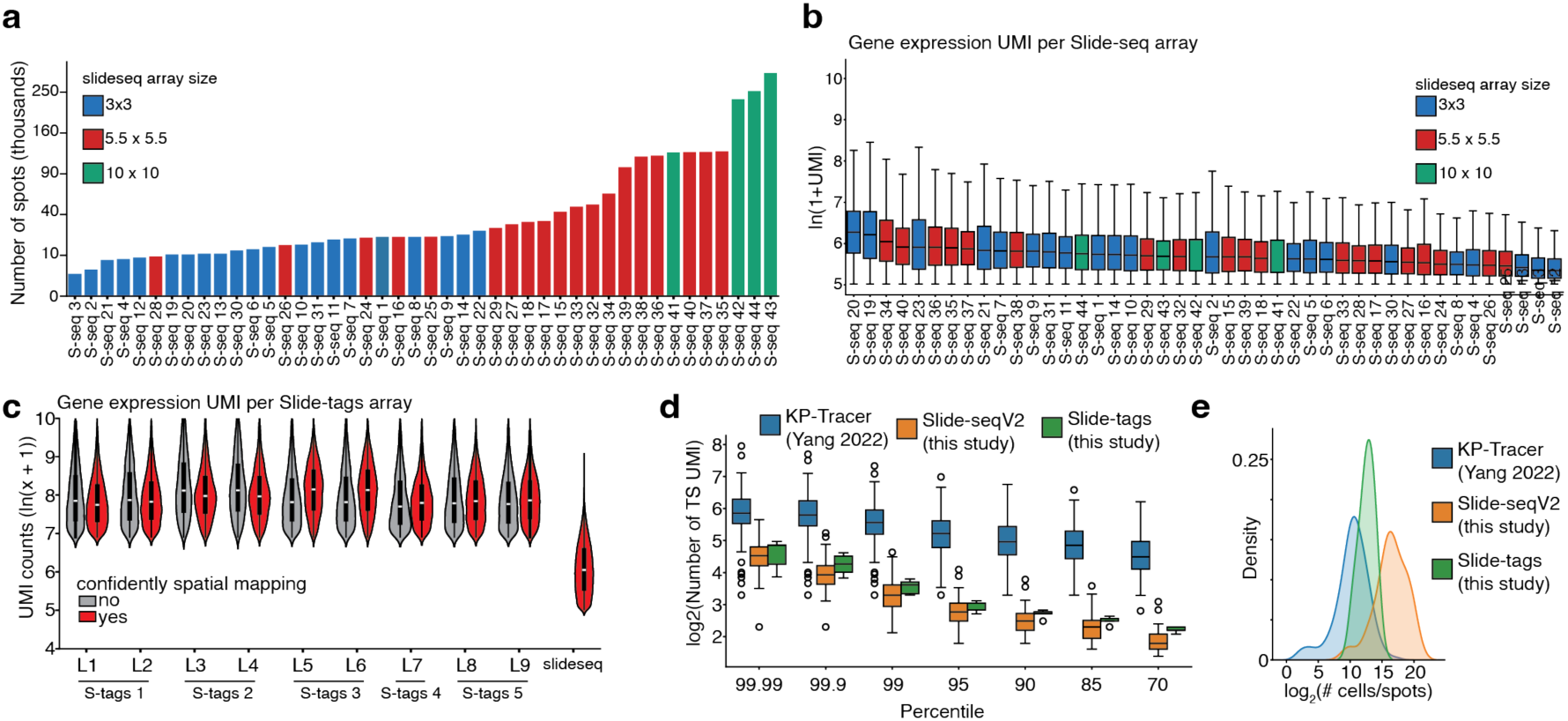
Characterization of spatial-lineage platform. **(a)** Number of spots that pass quality-control for all Slide-seq array. 3mm, 5mm, and 1cm arrays are uniquely coloured. **(b)** Number of gene expression UMIs for each Slide-seq array. Ln(1+UMI) is reported for each dataset. 3mm, 5mm, and 1cm arrays are uniquely coloured. Boxplots show the quartiles of the distribution, and whiskers extend to 1.5x the interquartile range. **(c)** Number of gene expression UMIs for each Slide-tags array, and one representative Slide-seq array. Each array is sequenced across multiple 10X libraries; assignment of 10X library to array is annotated. Distributions are split between cells that are confidently mapped and those that are not. Ln(1+UMI) is reported. **(d)** Distribution of number of target-site UMIs marking the top *X* percentile for whole-cell (KP-Tracer), Slide-seq, or Slide-tags datasets. Ln(1+UMI) is reported. Boxplots show the quartiles of the distribution, and whiskers extend to 1.5x the interquartile range. **(e)** Distribution of number of observations (cells or spots) that pass target-site quality-control in whole-cell (KP-Tracer), Slide-seq, or Slide-tags datasets. Log_2_ of the number of observations is reported.

**Extended Data Figure 2.**
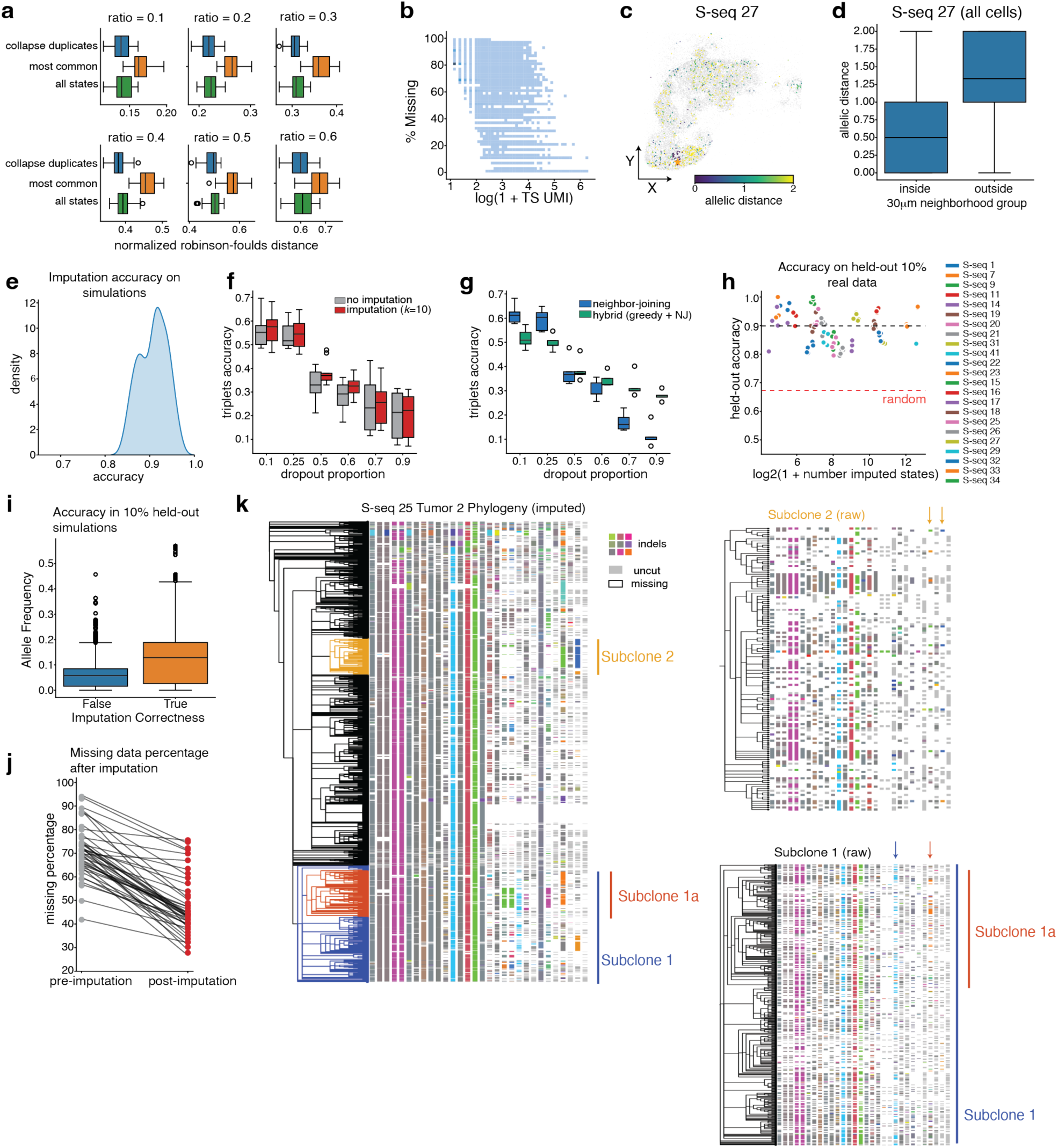
Benchmarking of computational approaches for spatial-lineage data. **(a)** Normalised Robinson-Foulds reconstruction error for simulated trees with increasing ratios of pooled cells and different pre-processing techniques. A ratio of *p* indicates that simulated lineage-tracing data of *p*% of cells are combined into a single observation to simulate multiple-cell capture in spatial transcriptomics (**Methods**). **(b)** Relationship between percentage of missing lineage-tracing data in a cell or spot and the log-number of UMIs (ln(1+x)) for Slide-seq and Slide-tags data. **(c)** Representative example of spatial coherence of lineage-tracing data on S-seq 27. For a selected spot (shown as a star), normalised allelic distance is reported for all spots with confident lineage-tracing data. Allelic distance is normalised between 0 and 2. **(d)** Distribution of allelic distances to spots within a 30𝜇𝑚 neighbourhood of a spot versus outside this neighbourhood. Distribution over all spots in S-seq 27 is reported. **(e)** Distribution of spatial imputation accuracy in lineage-tracing data simulated on a two-dimensional array. **(f)** Triplets-correct accuracy of reconstructed phylogenies simulated on a spatial array for various amounts of missing data rates, with and without spatial imputation. Boxplots show the quartiles of the distribution, and whiskers extend to 1.5x the interquartile range. **(g)** Triplets-correct accuracy of reconstructions with modified Neighbour-Joining and hybrid Cassiopeia-Greedy / Neighbour Joining algorithms for data simulated on a spatial array with various amounts of missing data, after spatial imputation. **(h)** Accuracy of spatial imputation and number of imputed states after holding-out 10% of all lineage-tracing data in Slide-seq datasets. Datasets where at least 10 imputations are made are shown. Median accuracy of random predictions is reported in a red dashed line. **(i)** Allele frequency of held-out data in a given tumour binned by imputation correctness. Boxplots show the quartiles of the distribution, and whiskers extend to 1.5x the interquartile range. **(j)** Overview of missing data reduction across all Slide-seq datasets after five rounds of spatial-imputation. **(k)** Phylogeny and lineage tracing heatmap of tree reconstructed in Fig. 1e. Subclones of interest are annotated in the same colours as in Fig. 1e. On the right, individual subclone heatmaps are presented with raw allelic data. Colour-coded arrows mark subclone-defining alleles. Unique colours of the heatmap indicate unique insertions or deletions (“indels”), white indicates missing data, and gray colours indicates no indel detected.

**Extended Data Figure 3.**
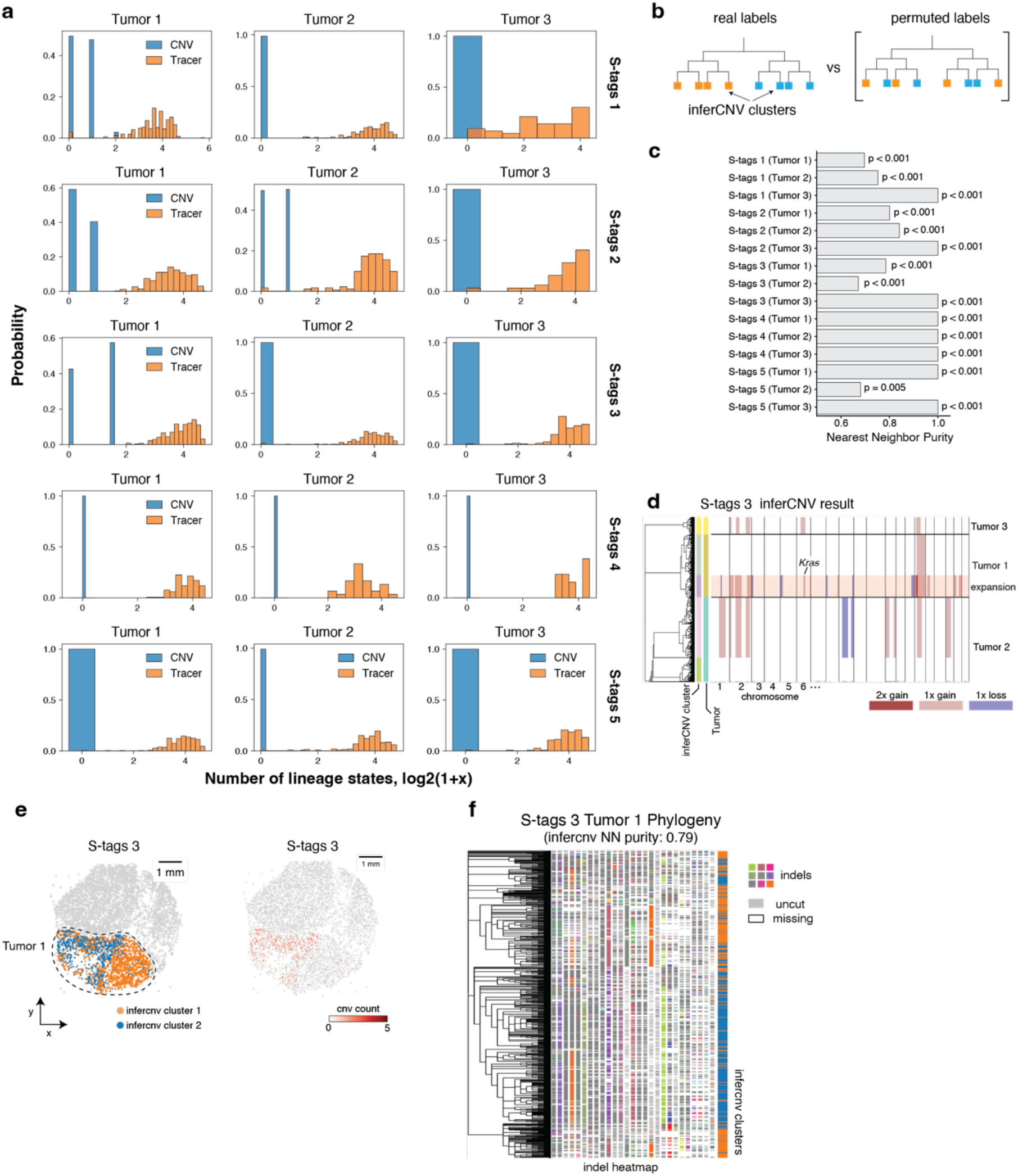
Characterization of copy-number variation in Slide-tags data. **(a)** Distribution of inferred CNV counts vs lineage-tracing states for each tumour in Slide-tags arrays. Tracer data consistently contains greater numbers of editable states; some tumours carry no inferred CNVs. **(b-c**) CNVs corroborate phylogenetic structure inferred from tracer data. (b) Schematic of nearest-neighbour test. Neighbours of each leaf in a phylogeny are tested for consistency in inferCNV cluster, and compared to a permutation null distribution. (c) Average nearest neighbour purity for each tumour, with empirical p-values computed against 1000 random permutations **(d-f)** inferCNV results for an example tumour. (d) inferCNV heatmap for a given Slide-tags dataset, S-tags 3. (e) CNV clusters and number of inferred CNVs per cell projected onto Tumour 1 in S-tags 3. (f) Tumour 1 phylogeny from S-tags 3, heatmap, and inferCNV cluster annotations.

**Extended Data Figure 4.**
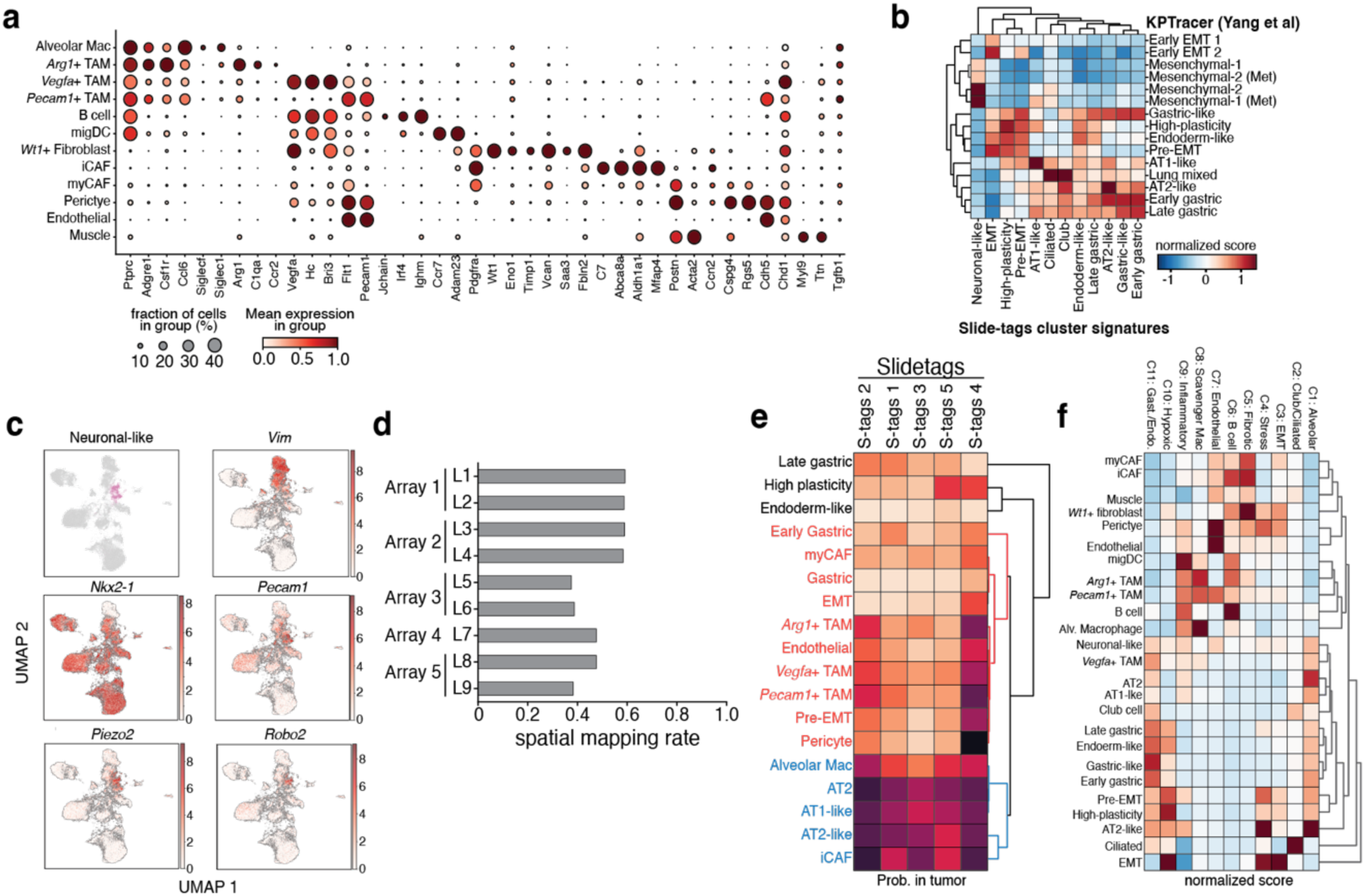
Analysis of Slide-tags data and cell-type annotation. **(a)** Summary of gene markers for each stromal cell population identified in Slide-tags. Each row corresponds to a stromal or immune cell-type cluster and each column corresponds to a marker gene. Dot size indicates the proportion of cells expression that gene, and colour indicates the average gene expression value (unit scaled between 0 and 1). **(b)** Clustered heatmap of transcriptional score of marker genes identified from Slide-tags data of tumour and epithelial cell types applied to previous KP-Tracer data. Scores are Z-normalised. **(c)** Annotation of Slide-tags tumour and epithelial UMAP projection with the Neuronal-like cell-type, and log-normalised gene expression patterns of selected genes: *Vim*, *Nkx2-1, Pecam1, Piezo2,* and *Robo2*. **(d)** Proportion of cells that are confidently mapped in each Slide-tags array. **(e)** Proportion of cells for each cell type that are found within the tumour boundary across Slide-tags arrays. **(f)** Clustered heatmap of transcriptional scores for each spatial community, identified from Hotspot analysis of Slide-seq data, for each Slide-tags cell type cluster. Scores are Z-normalised.

**Extended Data Figure 5.**
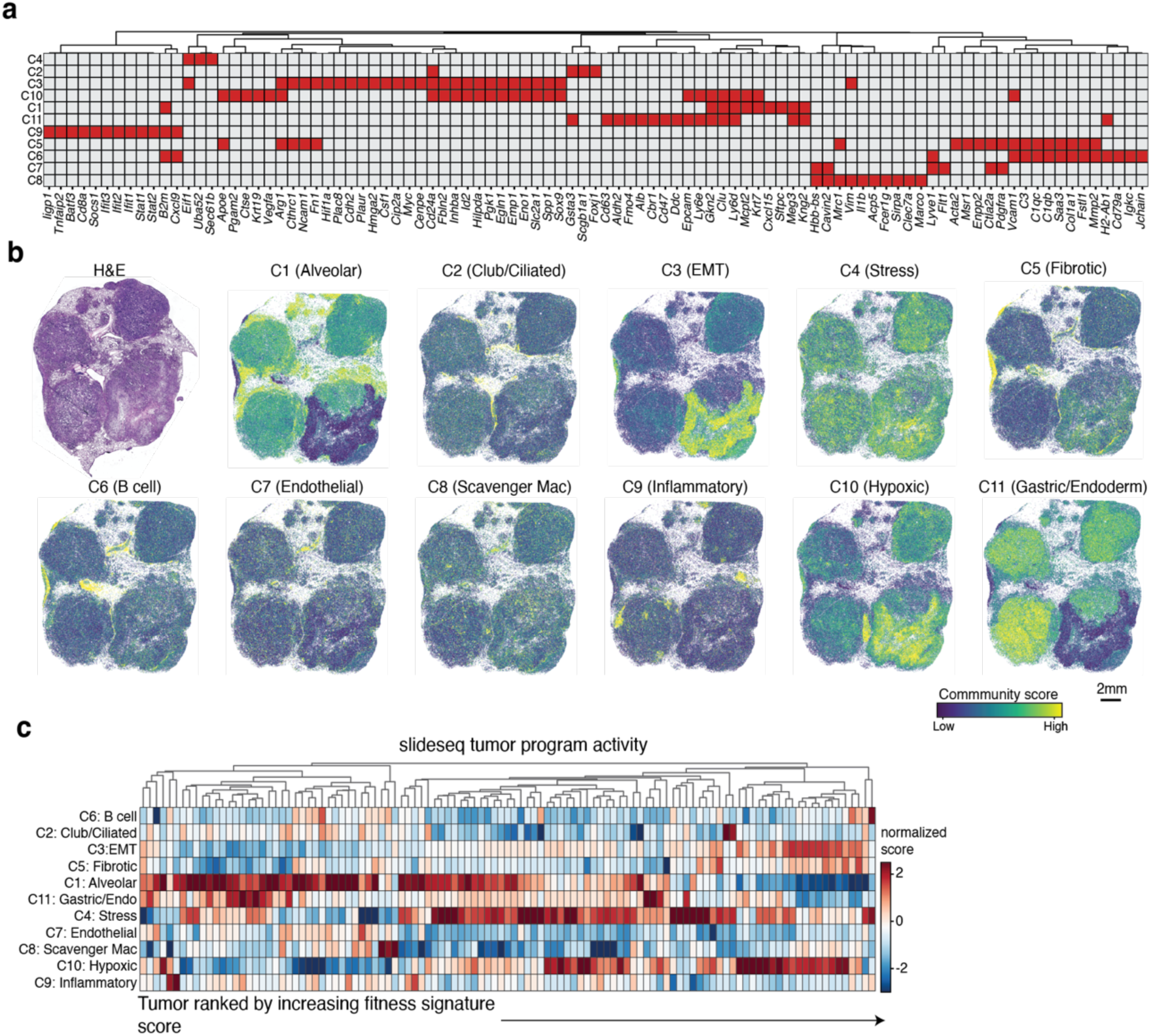
Analysis of spatial communities identified from Slide-seq data. **(a)** Clustered heatmap showing selected genes for each spatial community. Red colours indicate that a gene is found within that module. **(b)** Community scores for each spatial community and paired H&E for a representative Slide-seq community. **(c)** Clustered heatmap of community scores for each tumour in the Slide-seq dataset ordered by increasing fitness signature scores. Scores are Z-normalised.

**Extended Data Figure 6.**
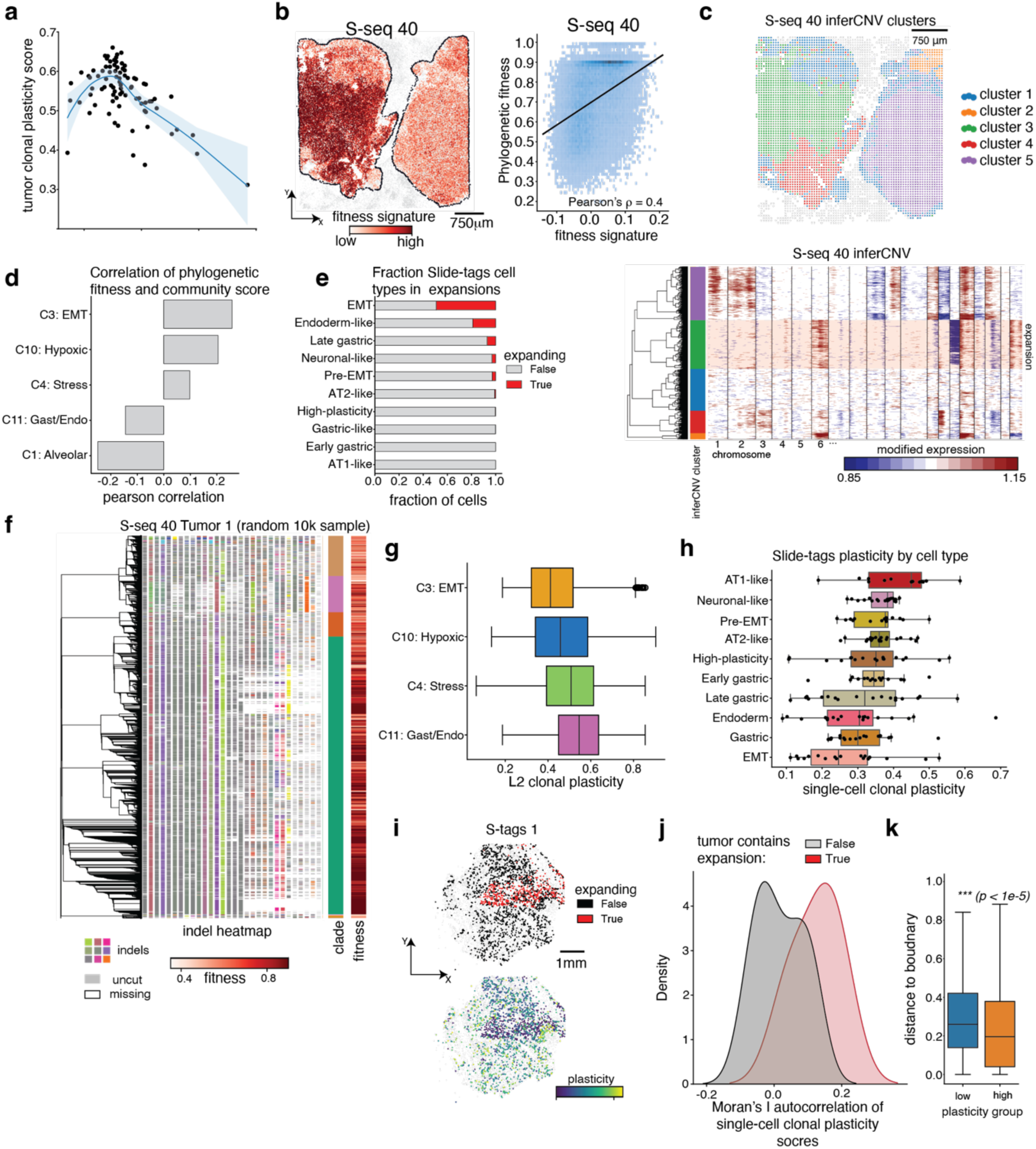
Characterization of subclonal tumour and microenvironmental dynamics. **(a)** Joint distribution of mean tumour clonal plasticity and fitness signatures across Slide-seq datasets. **(b)** Relationship between phylogenetic fitness, estimated from inferred trees, and transcriptional fitness signature score (Pearson’s correlation = 0.4) **(c)** Analysis of CNVs in example Slide-seq data, S-seq 40, shows a co-localization of a CNV clone with the hypoxic region of Tumour 1 shown in Figure 3a. The expanding CNV clone includes an amplification of a region on chr6 carrying *Kras*. **(d)** Correlation of phylogenetic fitness, estimated from inferred trees, and community scores for cancer-associated communities (C1: Alveolar; C3: EMT; C4: Stress; C10: Hypoxic; and C11: Gastric/Endoderm). Correlations are ordered in decreasing order. **(e)** Fraction of cells found in expanding regions of Slide-tags phylogenies, summarized for each cancer cell-type. **(f)** Reconstructed phylogeny and lineage tracing heatmap of representative tumour presented in Fig. 3a**-b**. Unique colours of the heatmap indicate unique insertions or deletions (“indels”), white indicates missing data, and gray colours indicates no indel detected. Colour bars indicate the subclonal clade and fitness, identical to those reported in Fig. 3a**-b**. **(g)** Distribution of L2 clonal plasticity (**Methods**) quantified in Slide-seq phylogenies summarized across spots annotated by cancer-dominated communities. **(h)** Distribution of single-cell clonal plasticity scores computed in Slide-tags phylogenies, stratified by cancer cell-types, and reported across tumour-array combinations. Boxplots show the quartiles of the distribution, and whiskers extend to 1.5x the interquartile range. **(i)** Representative spatial localization of phylogenetic expansion (top) and single-cell clonal plasticity scores (bottom) in a single Slide-tags array (S-tags 3). Scale bar indicates 1mm. **(j)** Distribution of autocorrelation values, computed by Moran’s I, of single-cell clonal plasticity scores for tumours with or without expansions. Higher autocorrelation values indicate that values have higher spatial coherence. Autocorrelations are reported across all Slide-tags datasets. **(k)** Distance to nearest non-tumour cell (i.e., tumour boundary) for high- and low-plasticity cells across all Slide-tags arrays. Cells with high-plasticity are closer to the tumour boundary (*p < 1e-5*, wilcoxon rank-sums test).

**Extended Data Figure 7.**
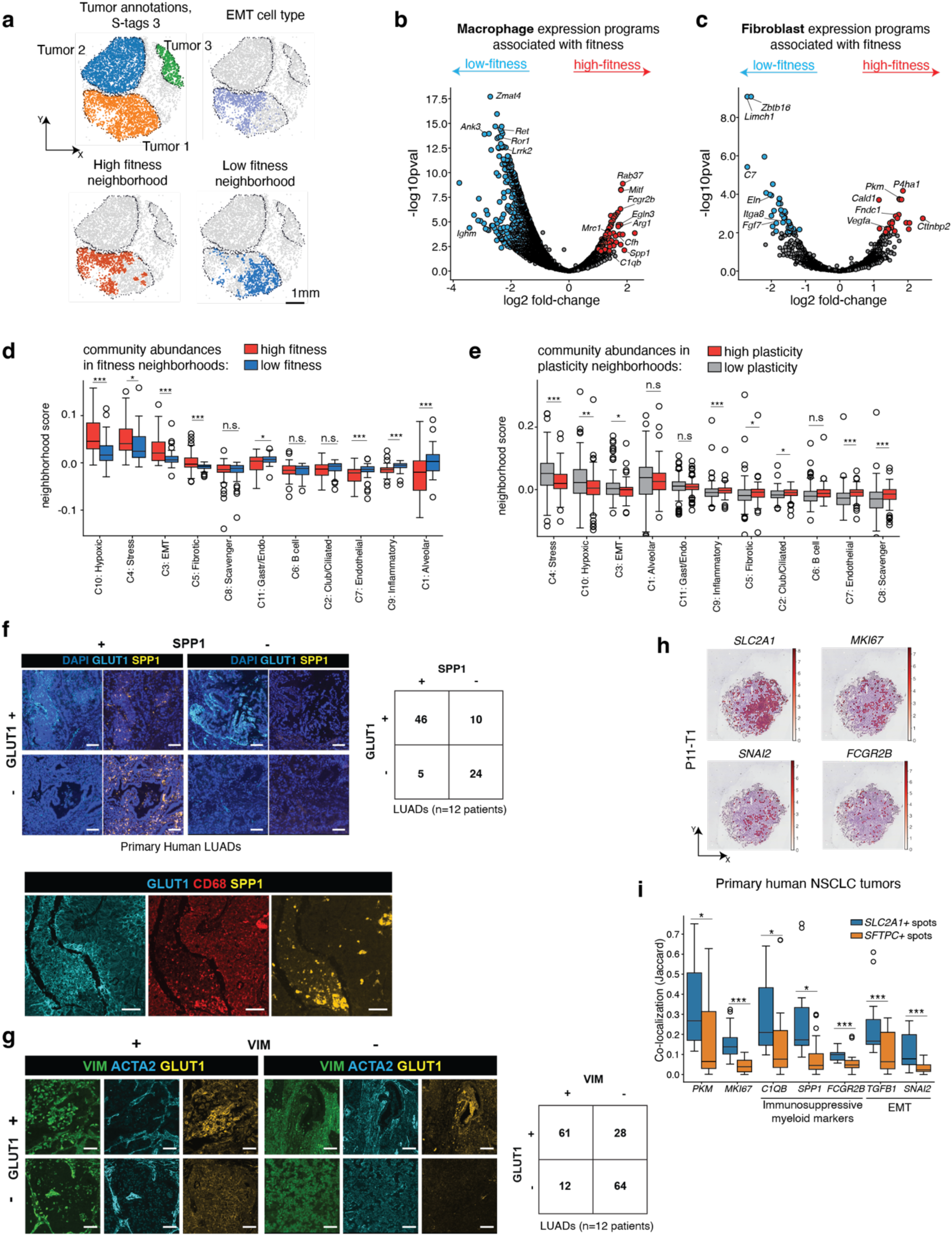
Analysis of cell-type-specific changes in the expanding tumor niche. **(a)** Representative example demonstrating the stratification of neighbourhoods of high- and low-fitness cells in Slide-tags data, and comparison to spatial localization of the EMT state. Scale bar indicates 1mm. **(b)** Differential expression analysis of macrophage polarisation states in neighbourhoods of high- and low-fitness cells from Slide-tags arrays. Each dot is a gene, and significant hits (log2|FC| >= 1 and false-discovery-rate adjusted p-value < 0.05) are reported in red and blue. Red genes are up-regulated in neighbourhoods of high-fitness cells, and blue genes are down-regulated. Significant GO terms are reported in **Supplementary Table 3**. **(c)** Differential expression analysis of fibroblast polarisation states in neighbourhoods of high- and low-fitness cells from Slide-tags arrays, as in **(b)**. **(d)** Distribution of average community scores in 30𝜇𝑚 neighbourhoods of high- or low-fitness spots in Slide-seq data. Each observation corresponds to a tumour. Significance is indicated above each comparison (*n.s.* = not significant; *\** = *p<0.1*; *\*\* = p<0.05; *** = p < 0.01*). Boxplots show the quartiles of the distribution, and whiskers extend to 1.5x the interquantile range. **(e)** Distribution of average community scores in 30 𝜇𝑚 neighbourhoods of high- or low-plasticity spots in Slide-seq data. Each observation corresponds to a tumour. Significance is indicated above each comparison (*n.s.* = not significant; *\** = *p<0.1*; *\*\* = p<0.05; *** = p < 0.01*). Boxplots show the quartiles of the distribution, and whiskers extend to 1.5x the interquantile range. **(f)** Immunofluorescence analysis of large lung adenocarcinoma surgical resections stained for SPP1, GLUT1, and CD68. Top: Representative images are shown (left) for a systematic quantification of co-localization between SPP1+ and GLUT1+ regions (right). GLUT1 & SPP1 are significantly co-localized (p<1e-5, Fisher’s Exact Test). Bottom: a representative image of CD68, GLUT1, and SPP1 co-localization. Scale bars are 100um. **(g)** Immunofluorescence analysis of large lung adenocarcinoma surgical resections stained for VIM, ACTA2, and GLUT. Representative images are shown (left) for a systematic quantification of co-localization between VIM+, ACTA2+, and GLUT1+ regions (right). VIM & GLUT1 are significantly co-localized (p<1e-5, Fisher’s Exact Test). Scale bars are 100um. **(h)** Representative example of spatial log-normalized gene expression values for selected genes in a human non-small-cell lung cancers (NSCLC) spatial transcriptomics dataset (see **Methods**). **(i)** Overall distribution of log-normalized gene expression values of selected genes co-expressed in hypoxic (*SLC2A1*+) or epithelial-like (SFTPC+) tumor spots across all NSCLC samples in dataset shown in (h). Ontologies are indicated underneath genes. Hypoxia+ spots (*SLC2A1*+) have higher expression of proliferation (*MKI67*), immunosuppressive myeloid (*FCGR2B*, *SPP1*, *C1QB*) and EMT (*SNAI2* and *TGFB1*) markers. Statistical significance between gene expression distributions is shown for each comparison (n.s. = not significant; * = *p*<0.1; ** = *p*<0.05; *** = *p* < 0.01; wilcoxon rank-sums test). Boxplots show the quartiles of the distribution, and whiskers extend to 1.5x the interquantile range.

**Extended Data Figure 8.**
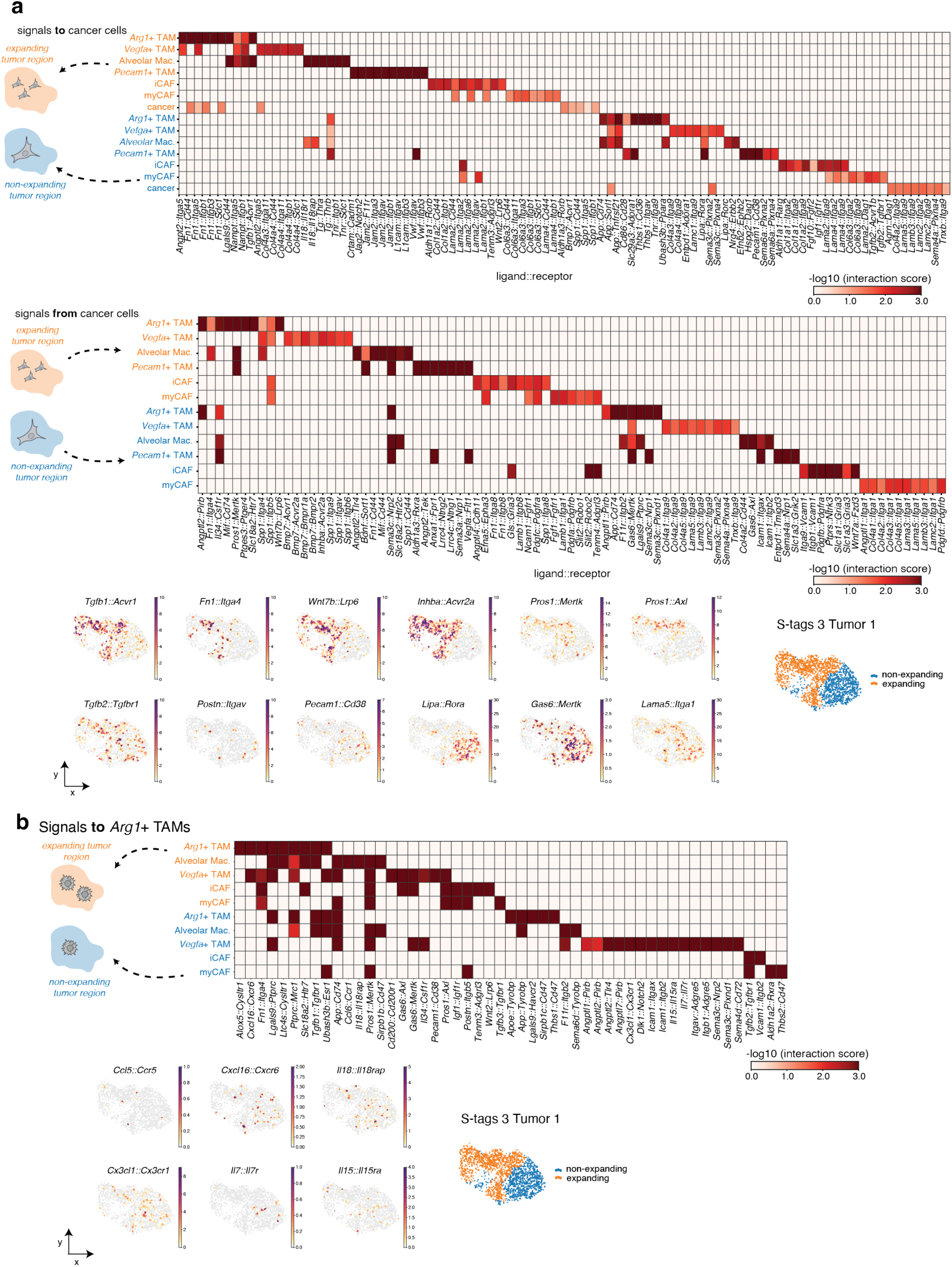
Spatially-aware, cell-type-specific ligand-receptor analysis in expanding niches. **(a)** Top ligand-receptor pairs are reported for cancer cells. Heatmaps indicate the average negative log10(interaction score) across 5 Slide-tags datasets, as computed with LARIS (see **Methods**). Representative spatial scatter plots are presented for specific ligand-receptor examples on S-tags 3 Tumour 1. **(b)** Top ligands for *Arg1+* TAMs in the expanding and non-expanding niches; heatmap and scatter plots are as above in **(a)**.

**Extended Data Figure 9.**
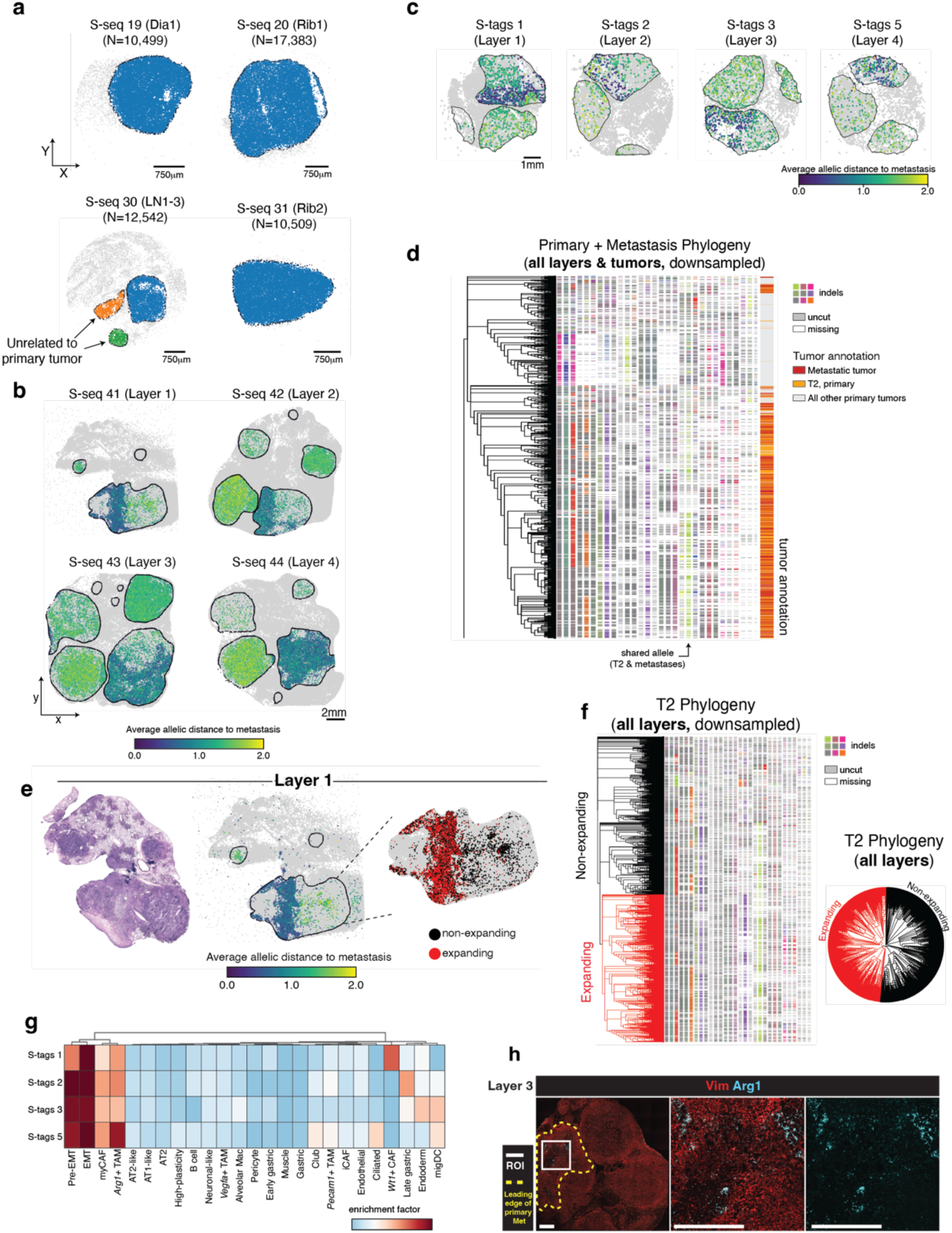
Profiling of step-wise transcriptional changes in the primary tumour during metastatic initiation. **(a)** Summary of metastases identified in Slide-seq spatial transcriptomics dataset. Each sample is annotated the metastatic site (LN: lymph node; Dia: Diaphragm). Two metastases in the lymph node (S-seq 30) were not found to be related to the primary tumour studies in Fig. 4 and thus removed from comparative analysis. **(b)** Spatial projection of allelic distances for each spot with lineage-tracing data to consensus metastatic parental allele across all four layers profiled in Slide-seq. Allelic distances are normalised between 0 and 2. **(c)** Spatial projection of allelic distances for each cell with lineage-tracing data to consensus metastatic parental allele across paired Slide-tags arrays. Allelic distances are normalised between 0 and 2. **(d)** Phylogeny and lineage tracing heatmap of lineage-tracing data from all tumours and metastases in analysis presented in Figure 4. To improve readability, each group is down sampled to at most 600 cells per group. Cells are annotated by group: metastatic tumours, T2 (primary tumour giving rise to metastases) or all other primary tumours in Slide-seq data. **(e)** H&E staining, spatial mapping of allelic distances to consensus metastatic parental allele state, and spatial localization of phylogenetic expansion for T2 in representative dataset. Allelic distances are normalised between 0 and 2. **(f)** Reconstructed phylogeny and lineage tracing heatmap of T2 from all layers. Unique colours of the heatmap indicate unique insertions or deletions (“indels”), white indicates missing data, and gray colours indicates no indel detected. Full circular phylogram is shown on the right. Clades participating in expansion shown in **(e)** are shown in red. **(g)** Clustered heatmap of enrichments of cell type abundances in spatial neighbourhoods of cells related to metastases in Slide-tags arrays. **(h)** Immunofluorescence imaging of ARG1 and VIM in a section of the tumour-bearing lung close to Layer 3. Leading edge of the metastasis-initiating subclone is indicated with yellow dashed line. Scale bar indicates 1mm.

**Extended Data Figure 10.**
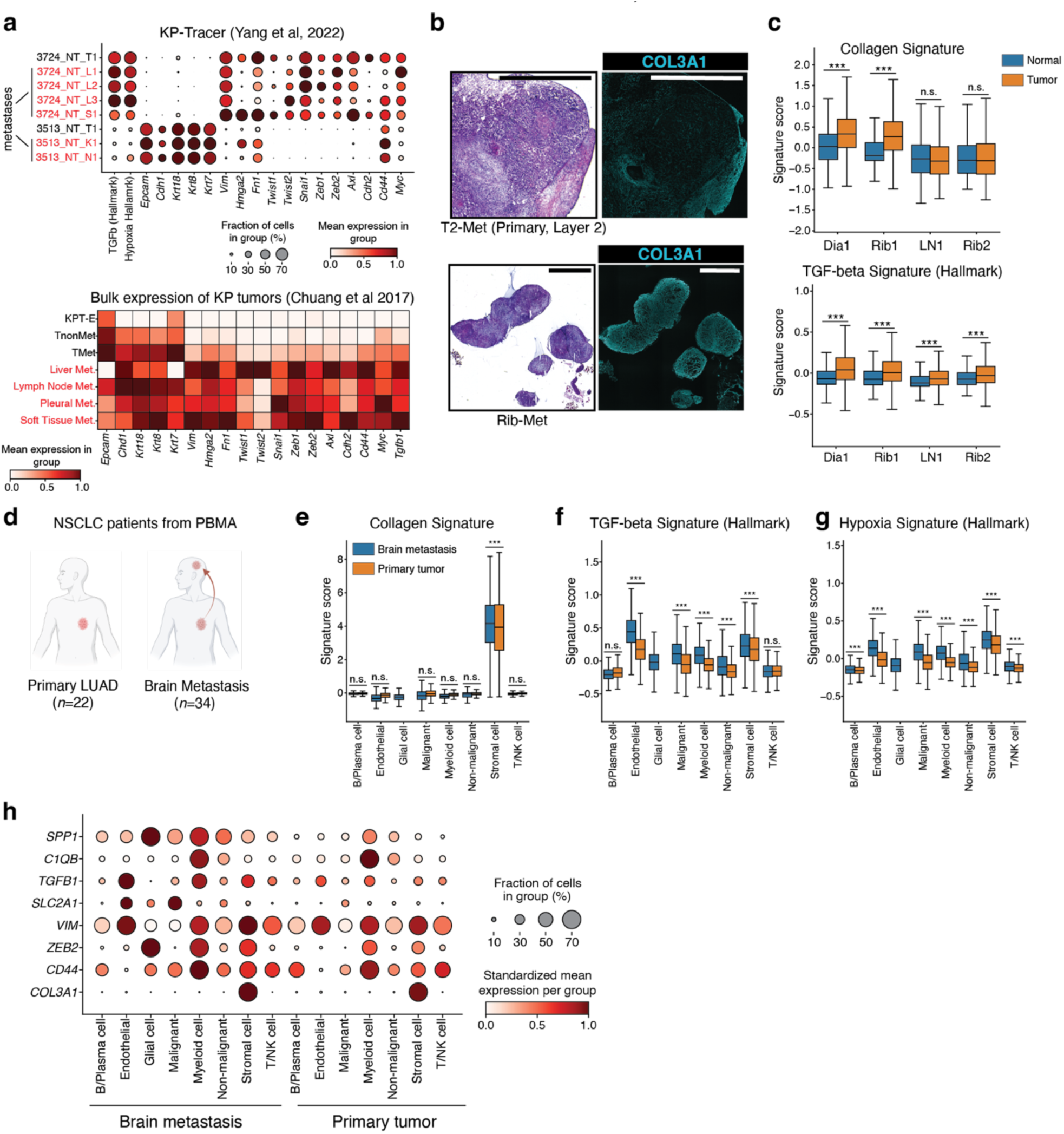
Analysis of microenvironmental changes associated with metastatic colonization. **(a)** Comparative analysis of metastasis-related gene expression between primary and metastatic tumours from KP tumours from the KP-Tracer study (top, *n*=6 metastases from 2 mice) and Chuang et al (bottom, *n*=24 metastases from 12 mice). Gene dots are sized by fraction of cell expressing a gene and values are standardized across genes (top). Metastases are indicated in red text. T: primary tumour, L: liver metastasis, S: soft tissue metastases, K: kidney metastasis, N: lymph-node metastasis; KPT-E: early stage KPT tumours; Tnonmet: primary tumors that do not seed metastases; Tmet: primary tumours that do seed metastases. **(b)** H&E and immunofluorescence imaging of COL3A1 in a section of the metastasis-initiating primary tumour (Layer 2) and related metastasis. Scale bar indicates 2.5mm. **(c)** Expression of collagen deposition and *TGF* 𝛽 signalling pathways in Slide-seq data, segmented by tumour region. Significant increases in tumour region are indicated (***: p<0.01; n.s.: not significant). **(d)** Analysis of 56 human lung adenocarcinoma tumours from the PBMA cohort, including 34 brain metastases and 22 primary LUAD tumours from the PBMA LUAD cohort. **(e-g)** Cell-type specific transcriptional signatures of collagen deposition, *TGF*𝛽 signalling, and hypoxia across compartments (brain metastasis or primary tumour). Significant increases in brain metastases are indicated (***: p<1e-3; n.s.: not significant). **(h**) Expression of specific marker genes across cell types and compartments. Dots are sized by fraction of cells expressing the gene, and expression values are standardized across genes.

